# *Medicago truncatula* Ferroportin2 mediates iron import into nodule symbiosomes

**DOI:** 10.1101/630699

**Authors:** Viviana Escudero, Isidro Abreu, Manuel Tejada-Jiménez, Elena Rosa-Núñez, Julia Quintana, Rosa Isabel Prieto, Camille Larue, Jianqi Wen, Julie Villanova, Kirankumar S. Mysore, José M. Argüello, Hiram Castillo-Michel, Juan Imperial, Manuel González-Guerrero

## Abstract

Iron is an essential cofactor for symbiotic nitrogen fixation. It is required by many of the enzymes facilitating the conversion of N_2_ into NH_4_^+^ by endosymbiotic bacteria living within root nodule cells, including signal transduction proteins, O_2_ homeostasis systems, and nitrogenase itself. Consequently, host plants have developed a transport network to deliver essential iron to nitrogen-fixing nodule cells. Model legume *Medicago truncatula Ferroportin2* (*MtFPN2*) is a nodule-specific gene that encodes an iron-efflux protein. MtFPN2 is located in intracellular membranes in the nodule vasculature, and in the symbiosome membranes that contain the nitrogen-fixing bacteria in the differentiation and early-fixation zones of the nodules. Loss-of-function of *MtFPN2* leads to altered iron distribution and speciation in nodules, which causes a reduction in nitrogenase activity and in biomass production. Using promoters with different tissular activity to drive *MtFPN2* expression in *MtFPN2* mutants, we determined that MtFPN2-facilitated iron delivery across symbiosomes is essential for symbiotic nitrogen fixation, while its presence in the vasculature does not seem to play a major role in in the conditions tested.

## INTRODUCTION

Iron is an essential cofactor for enzymes participating in almost every plant physiological process, including photosynthesis, respiration, and stress tolerance (Marschner, 2011). Consequently, when lacking, a plethora of iron-dependent processes is affected. Damaging as low iron levels are, so are slightly higher ones. In that situation, iron becomes toxic, catalysing the production of free radicals in Fenton-style reactions and competing with other essential metal nutrients for the active site of proteins (Goldstein et al., 1993; Lin et al., 2011). To maintain iron homeostasis within these narrow physiological levels, plants have developed a complex system to optimize iron uptake and delivery from soil, while preventing toxicity. This system involves metal transporters that recover this nutrient from soil and deliver it to shoots and seeds (Korshunova et al., 1999; Curie et al., 2001; Vert et al., 2002; Morrissey et al., 2009; Zhai et al., 2014), small soluble molecules that coordinate iron and keep it soluble in plant fluids (von Wiren et al., 1999; Rogers and Guerinot, 2002; Roschzttardtz et al., 2011; Flis et al., 2016), and a complex regulatory network (Colangelo and Guerinot, 2004; Long et al., 2010; Sivitz et al., 2012; Kumar et al., 2017; Grillet et al., 2018). While in most vegetative growing plants, the main iron sinks are the leaves, legumes may have additional ones in differentiated root organs known as nodules (Brear et al., 2013; González-Guerrero et al., 2014). Representing just 5% of the total biomass of the plant, nodules contain a large part of the plant iron content (González-Guerrero et al., 2014; Tejada-Jiménez et al., 2015). Therefore, it is to be expected that legumes have adapted their pre-existing iron homeostasis systems to accommodate nodule development and function.

Nodules are the result of the interaction between legumes and rhizobia, a diverse group of soil bacteria capable of converting, fixing, N_2_ into NH_4_^+^ when in symbiosis (Downie, 2014). Following a complex signal exchange between the symbionts (Vernié et al., 2015), nodule primordia emerge from the roots by proliferation of pericycle and inner cortex cells (Xiao et al., 2014). While the nodule develops, rhizobia in the root surface are surrounded by a root hair that invaginates and directs, via an infection thread, the bacteria from the root exterior to the inner nodule regions (Gage, 2002), where they are released in an endocytic-like process (Limpens et al., 2009; Huisman et al., 2012). Within these pseudo-organelles (symbiosomes), rhizobia differentiate into bacteroids (Sutton et al., 1981; Kereszt et al., 2011), and start fixing nitrogen, transferring fixed NH_3_ to the host plant in exchange for photosynthates and mineral nutrients (Udvardi and Poole, 2013). Nodules can be classified according to their development in determinate and indeterminate (Sprent, 2007). The former, as illustrated by soybean nodules, has a limited growth, while the later (developed by pea or *Medicago*), maintains the apical meristem and has a longer growth period. Due to this indeterminate nodule growth, four nodule zones are established, tracing along its length the developmental stages of nodule maturation. These zones are: the meristem (zone I), the zone where the nodule cells are infected by rhizobia and they differentiate into bacteroids (zone II), the N_2_ fixation zone (zone III), and in nodules of a certain age, the senescent zone (zone IV) (Vasse et al., 1990). In addition, some authors also differentiate an interzone between zones II and III in which O_2_ levels drop and nitrogenase is starting to be synthesized (Roux et al., 2014).

Nodule development is highly sensitive to iron levels, as suggested by its abundance in these organs. Iron deficiency limits nodule number and maturation, as well as reduces nitrogen fixation rates (O’Hara et al., 1988; Tang et al., 1990; Tang et al., 1992). This is the consequence of many of the key enzymes involved in nodulation and nitrogen fixation being ferroproteins synthesized at high levels (Brear et al., 2013; González-Guerrero et al., 2014). For instance, heme-containing NADPH-oxidases are important for nodule initiation and development (Montiel et al., 2012; Montiel et al., 2016). Nitrogenase, the critical enzyme for nitrogen fixation, is a protein complex with 34 iron atoms in the form of iron-sulfur (Fe-S) clusters and the unique iron-molybdenum cofactor (FeMoco) (Rubio and Ludden, 2005). This enzyme is irreversibly inhibited by O_2_, while bacteroids must carry out an aerobic metabolism (Preisig et al., 1996b). To make both processes compatible, legumes express leghemoglobin (20% of the total nodule protein) which binds O_2_ with great affinity (Appleby, 1984). This oxygen can be recovered and used by high-affinity rhizobial cytochrome *c* oxidases (iron-copper proteins) (Preisig et al., 1996a; Preisig et al., 1996b). In addition, a number of other iron-proteins are required for free radical control, cofactor assembly, and general metabolic processes (Dalton et al., 1998; Rubio et al., 2004; Rubio and Ludden, 2005; González-Guerrero et al., 2014).

To ensure the reinforced iron supply that symbiotic nitrogen fixation requires, nodulation elicits the iron-deficiency response characteristic of dicots (Terry et al., 1991): lowering the pH of the rhizosphere to increase Fe^3+^ solubility, activating ferroreductase activities to reduce Fe^3+^ to Fe^2+^, and inducing the expression of Fe^2+^ importers of the ZIP (Zrt1-, Irt1-like Proteins) and Nramp (Natural Resistance Associated Macrophage Protein) transporter families (Andaluz et al., 2009; Kobayashi and Nishizawa, 2012). Once iron is in the plant, it symplastically or apoplastically reaches the endodermis from where it will be delivered by the vasculature to shoots and nodules. At the nodules, iron is released in the apoplast of the infection zone of the nodule (Rodríguez-Haas et al., 2013), where it is kept soluble as iron-citrate (Kryvoruchko et al., 2018), the most abundant complex in root to shoot iron transport and when transferring it across symplastically disconnected tissues (Durrett et al., 2007; Roschzttardtz et al., 2011). Iron uptake into the infected cells of *Medicago truncatula* is mediated primarily by MtNramp1, a transporter expressed in late zone II and interzone and located in the plasma membrane (Tejada-Jiménez et al., 2015). Transport into the symbiosomes is likely also facilitated by citrate, as indicated by the requirement of citrate efflux protein MtMATE67 (Kryvoruchko et al., 2018). However, this protein does not transport iron-citrate complexes, and therefore, another system must exist for iron translocation. Different candidates have been proposed to be responsible for iron delivery into the symbiosomes. The first was *Glycine max* Nramp protein GmDMT1 (Kaiser et al., 2003). However, although this protein was located in symbiosomes, its direction of transport would be into the cytosol, as with all characterized Nramp proteins (Nevo and Nelson, 2006). VIT1-CCC1-like protein SEN1 is another candidate to carry out this role (Hakoyama et al., 2012), but although the direction in which iron is transported by these proteins would be compatible (Li et al., 2001), there is no definitive evidence of a symbiosome localization. Alternatively, a member of the ferroportin family (also known as IREG proteins) could also be involved.

Ferroportins are membrane proteins that extrude iron from the cytosol, either outside the cell or into organelle (Morrissey et al., 2009; Drakesmith et al., 2015). In plants, ferroportins have been associated to iron delivery into the xylem (Morrissey et al., 2009). Interestingly, this is a process that would be coupled to MATE-mediated citrate efflux (FRD3 in *A. thaliana*) (Durrett et al., 2007), as iron is kept soluble in the xylem as an iron-citrate complex (López-Millán et al., 2000; Durrett et al., 2007). Considering the functional association of ferroportins and citrate efflux proteins in the xylem, we hypothesized that a similar relationship would be established between nodule-specific citrate efflux protein MtMATE67 and a ferroportin to deliver iron to nodule apoplast and to symbiosomes. As a result, we have identified and characterized MtFPN2 (*Medtr4g013240*), one of the three ferroportin-like genes encoded in the *M. truncatula* genome, and the only one with a nodule-specific expression. MtFPN2 is located in the intracellular compartments in the endodermis, and associated to symbiosomes in the infected cells. Mutation of *MtFPN2* results in a severe reduction of nitrogenase activity, and in mis-localization of iron in the nodules. These results support a role of MtFPN2 in iron delivery for symbiotic nitrogen fixation.

## RESULTS

### *MtFPN2* is expressed in the nodule vasculature and interzone

Three ferroportin-like sequences could be found in the genome of *M. truncatula*: *Medtr1g084140* (*MtFPN1*), *Medtr4g013240* (*MtFPN2*), and *Medtr4g013245* (*MtFPN3*). Among them, *MtFPN2* was the only ferroportin to have a nodule-specific transcription in the conditions tested (Figure 1A; Supplemental Figure 1). To determine the area of the nodule where *MtFPN2* was expressed, the 2,144 bp DNA region immediately upstream of *MtFPN2* start codon was used to drive the expression of the reporter ***β***-*glucuronidase* (*gus*) gene. After 28 days-post-inoculation (dpi), GUS activity was detected in a specific nodule band close to the apical region, and in longitudinal bands along its major axis (Figure 1B). Longitudinal sections of nodules showed that the apical GUS-activity region corresponded to the late infection zone, interzone and early fixation zone (Figure 1C). Nodule cross sections showed GUS activity in the nodule vasculature (Figure 1D). In these cross sections, the lack of expression of *MtFPN2* in the matured fixation zone was also confirmed.

**Figure 1.**
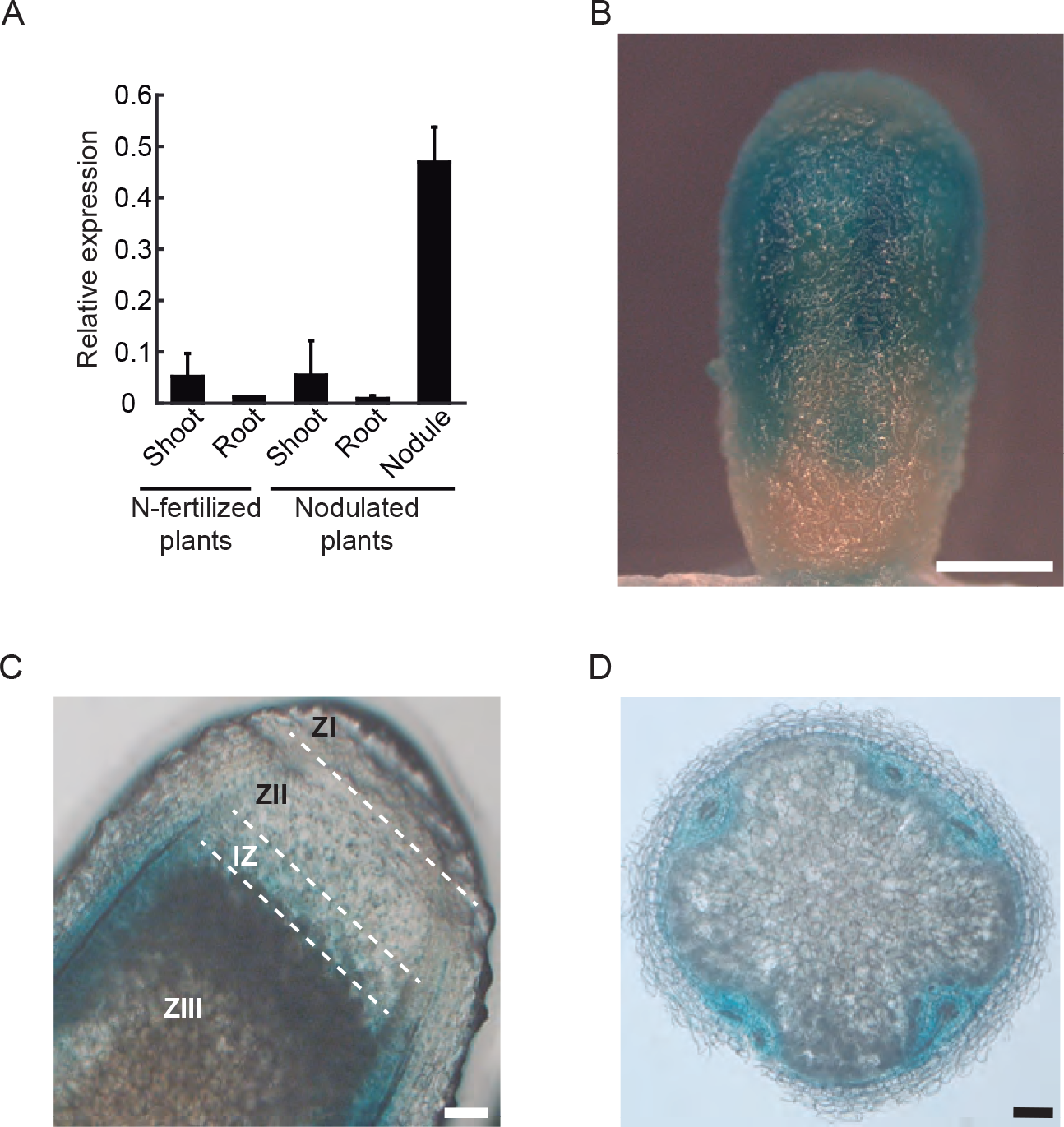
*MtFPN2* is expressed in the vasculature and in the differentiation-fixation zones of *M. truncatula* nodules. (A) *MtFPN2* expression in nodulated or nitrogen-fertilized *M. truncatula* plants relative to the internal standard gene *Ubiquitin carboxyl-terminal hydrolase*. Data are the mean ± SE of two independent experiments with 4 pooled plants. (B) GUS staining of 28 dpi *M. truncatula* nodules expressing the *gus* gene under the control of *MtFPN2* promoter region. Bar = 500 μm. (C) Longitudinal section of a GUS-stained 28 dpi *M. truncatula* nodule expressing the *gus* gene under the *MtFPN2* promoter region. ZI indicates Zone I; ZII, Zone II; IZ, Interzone; and ZIII, Zone III. Bar = 100 μm. (D) Cross section of a GUS-stained 28 dpi *M. truncatula* nodule expressing the *gus* gene under the *MtFPN2* promoter region. Bar = 100 μm.

### MtFPN2-HA is located in vascular intracellular compartments and in the symbiosomes

Ferroportins have been associated to iron efflux from the cytosol (Kaplan, 2002). Their specific functional role is largely derived from the specific subcellular localization of each protein. Should it be associated to the plasma membrane, its likely role would be iron export for long-distance trafficking, such as FPN1 in enterocytes, or FPN1 in endodermal cells (McKie et al., 2000; Morrissey et al., 2009). Alternatively, an intracellular localization associated to organelle would mean a role in iron storage or metalation of specific ferroproteins (Morrissey et al., 2009; Drakesmith et al., 2015). To determine the subcellular distribution of MtFPN2 and gain insights into its function, three hemagglutinin (HA) epitopes where fused to its C-terminus. The expression of this cassette was driven by the same region previously used for the *MtFPN2*_prom_∷*gus* fusions. Protein localization was initially determined by confocal microscopy in which the epitopes were detected by an anti-HA monoclonal primary antibody and an Alexa594-conjugated (red) secondary antibody. To facilitate visualization, the transformed plants were inoculated with a GFP-expressing *Sinorhizobium meliloti* strain (green) and the sections stained with DAPI (blue) to visualize DNA. Figure 2A shows that MtFPN2-HA localization corresponded to the expression profile detected with the *MtFPN2*_prom_∷*gus* fusions. A detailed view of the vascular region revealed that MtFPN2-HA was located in the endodermis, pericycle and in the vascular parenchyma, occupying large areas of the cytosol (Figure 2B). Visualization of infected cells in the differentiation zone also showed an intracellular localization of MtFPN2-HA, closely related to the symbiosomes (Figure 2C). To confirm this putative symbiosome localization, *M. truncatula* plants were transformed with a C-terminal GFP-fusion of *MtFPN2* driven by its own promoter and inoculated with a mCherry expressing *S. meliloti*. Symbiosomes isolated from these nodules showed co-localization of rhizobia and MtFPN2-GFP (Supplemental Figure 2). To further verify the symbiosome localization of MtFPN2-HA, a gold-conjugated secondary antibody was used to determine the distribution of HA-tagged proteins using transmission electron microscopy. As indicated by the confocal microscopy assays, gold particles were associated to symbiosome membranes (Figure 2D). The same approach showed gold particles concentrated in large intracellular membrane systems, very likely the endoplasmic reticulum in endodermal and vascular parenchyma cells (Figure 2D). To confirm that none of these observations were artifactual, controls for autofluorescence of the tagged proteins and specificity of the antibodies were carried out (Supplemental Figure 3).

**Figure 2.**
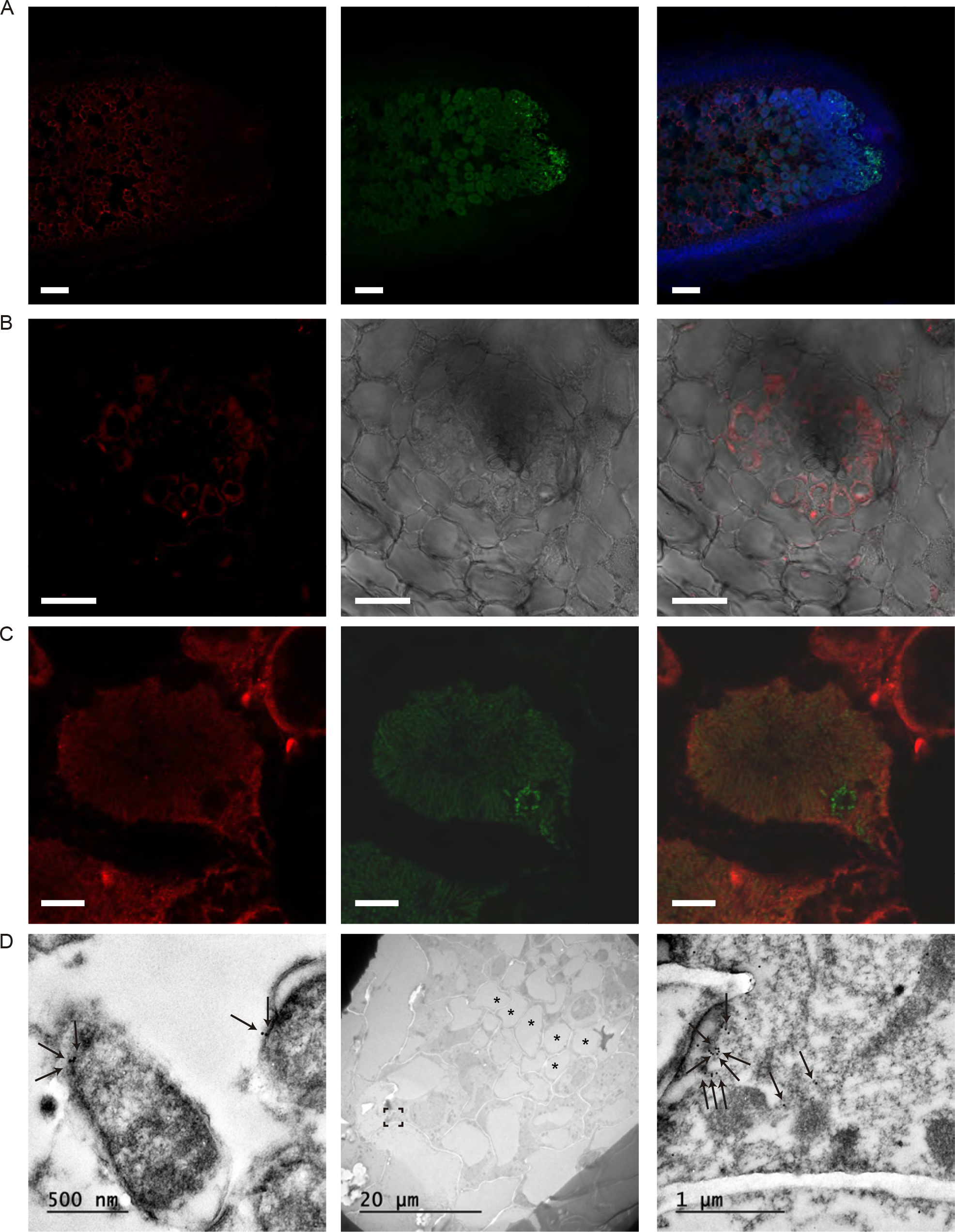
MtFPN2 is localized in intracellular membranes in the vasculature and in the symbiosome membrane in *S. meliloti*-infected nodule cells. (A) Longitudinal section of a 28 dpi *M. truncatula* nodule expressing *MtFPN2-HA* under its own promoter. The three C-terminal HA epitopes were detected using an Alexa594-conjugated antibody (red, left panel). Transformed plants were inoculated with a GFP-expressing *S. meliloti* strain (green, middle panel). Both images were overlaid with the channel showing DAPI-stained DNA (blue, right panel). Bar = 100 μm. (B) Cross section of a vessel of 28 dpi *M. truncatula* nodule expressing *MtFPN2-HA* under its own promoter. The three C-terminal HA epitopes were detected using an Alexa594-conjugated antibody (red, left panel). Transillumination image is shown in the centre panel, and overlaid with the Alexa594 signal (right panel). Bar = 25 μm. (C) Detail of a *S. meliloti*-colonized 28 dpi nodule cell in the differentiation zone. Right panel shows the Alexa594 signal (red), middle panel the *S. meliloti* distribution (green), and the right one overlays both images. Bar = 10 μm. (D). Detail of symbiosomes shown by transmission electron microscopy (left panel). Arrows indicate the position of the colloidal gold particles conjugated to the antibody used to detect MtFPN2-HA. Middle panel, *M. truncatula* nodule vessel shown with transmission electron microscopy. Asterisks indicate the position of the xylem. Boxed region was amplified in the right panel to show the antibody-conjugated gold particles used to position MtFPN2-HA (indicated by arrows).

### MtFPN2 is an iron efflux protein

To confirm that MtFPN2 was capable of transporting iron, we tested its ability to restore the wild-type growth of a yeast mutant lacking the CCC1 protein. This transporter introduces excess cytosolic iron into the vacuole to prevent toxicity (Li et al., 2001). Loss of CCC1 function resulted in lower iron tolerance when the media was supplemented with 2.5 mM FeSO_4_. However, when this strain was transformed with a plasmid expressing the coding sequence of *MtFPN2*, the sensitivity of the mutant strain to high iron was diminished (Figure 3), suggesting that MtFPN2 was able to participate in its detoxification. Addition of the HA-tag used for the immunolocalization studies did not affect MtFPN2 activity in yeast (Supplemental Figure 4).

**Figure 3.**
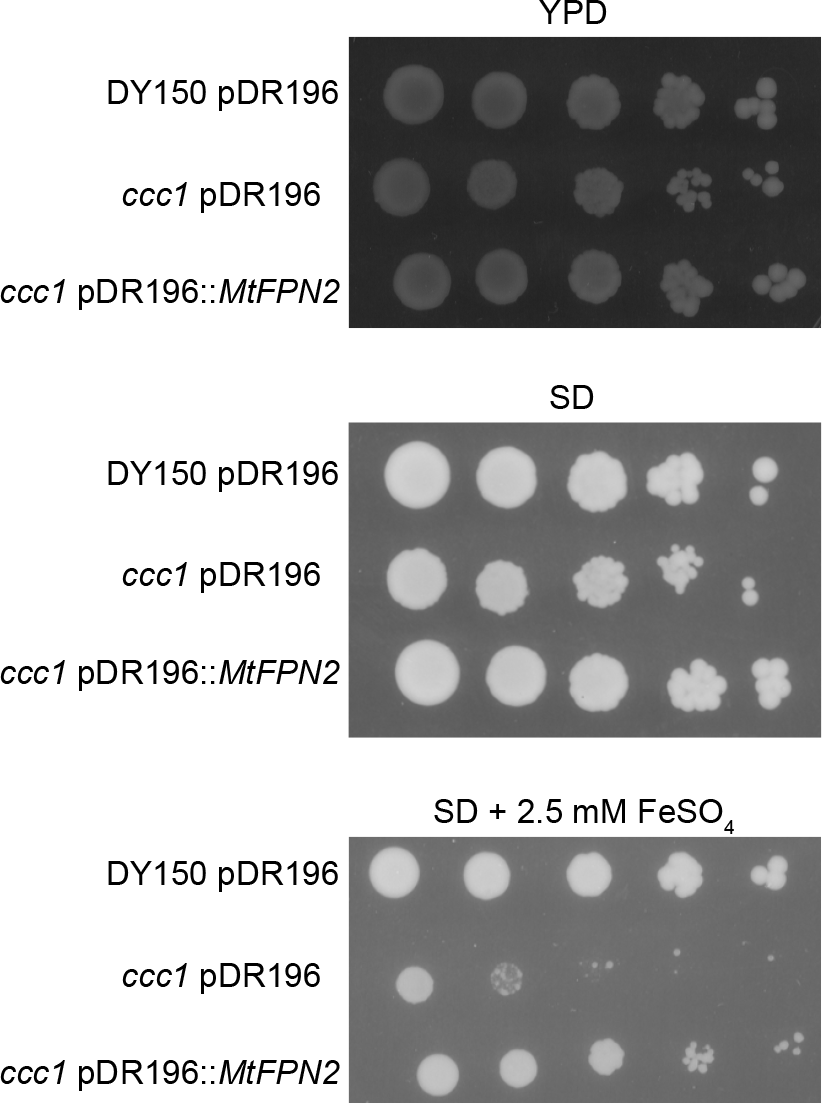
MtFPN2 transports iron out of the cytosol. Parental yeast strain DY150 was transformed with the pDR196 empty vector, while the *ccc1* mutant was transformed with either pDR196 or pDR196 harbouring the *MtFPN2* coding sequence. Serial dilutions (10x) of the transformants were grown in YPD medium (top panel), SD medium with the required auxotrophies (centre panel), or in SD medium with the required auxotrophies and 2.5 mM FeSO_4_ (lower panel).

### MtFPN2 is required for nitrogenase activity in nodules

Should MtFPN2 be responsible for iron delivery to the nodules, its mutation should lead to a reduction of nitrogenase activity, as a consequence of an iron-limited cofactor synthesis. To explore this possibility, the *Tnt1* insertion mutant NF11374 (*fpn2-1*) was obtained from the Noble Research Institute (Tadege et al., 2008). This line has an insertion in the fourth exon of *MtFPN2* (Figure 4A), resulting in no transcript being detected by RT-PCR. *MtFPN2* loss-of-function had no significant effect on plant growth or chlorophyll content when the plants had not been inoculated and nitrogen was provided as ammonium nitrate in the nutrient solution (Supplemental Figure 5). However, when the plants were inoculated and nitrogen had to be obtained from the endosymbiotic rhizobia, removing *MtFPN2* function led to severe growth reduction (Figure 4B). Nodule development was also altered, with a large portion (64%) of the nodules having white colour (Figure 4C, 4D), indicative of non-functionality. Closer observation of these nodules using either toluidine blue-stained sections or tomographic reconstructions from X-ray analyses, showed that the white *fpn2-1* nodules did not develop the fixation zones further than a few cell layers (Supplemental Figure 6), and that even the fewer nodules with a pink-colour had a reduced Zone III. In addition, the symbiosomes appeared disorganized in the cytosol of *fpn2-1* nodule cells, with a number of vacuoles, in contrast to the ordered, radial and peripheral distribution in the wild type plants. As expected by the plant and nodule phenotypes, biomass production was significantly lower in *fpn2-1* plants than in wild-type plants (Figure 4E). Nitrogen fixation capabilities (nitrogenase activity) of the mutant nodules were approximately 80% lower than those from wild-type plants (Figure 4F). These phenotypes were the likely result of iron not reaching the nodules in high-enough levels, as indicated by the lower iron content of *fpn2-1* nodules (Figure 4G). Introducing into *fpn2-1* the coding sequence of *MtFPN2* regulated by the 2,144 bp region upstream its start codon restored the wild-type phenotype. However, fortifying the nutrient solution with additional iron did not have the same effect (Supplemental Figure 7).

**Figure 4.**
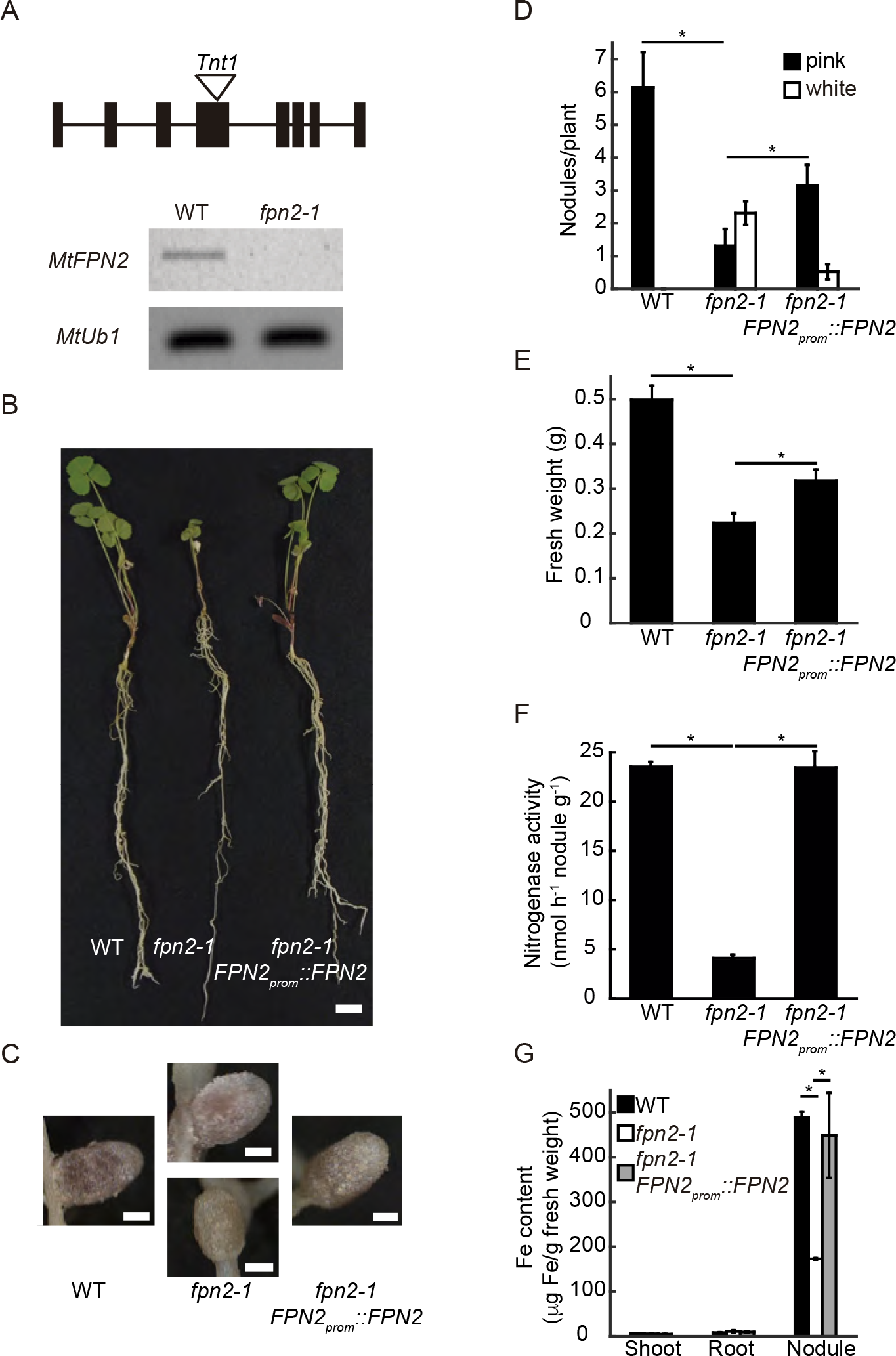
MtFPN2 is required for symbiotic nitrogen fixation. (A) *Tnt1* insertion in the fourth exon of *MtFPN2* results in no expression being detected by RT-PCR. The *Ubiquitin carboxyl-terminal hydrolase* (MtUb1) gene was used as a positive control. (B) Growth of representative wild type (WT), *fpn2-1*, and *fpn2-1* transformed with the *MtFPN2* coding sequence expressed under its own promoter (*FPN2*_*prom*_∷*MtFPN2*). Bar = 1 cm. (C) Detail of representative nodules of WT, *fpn2-1*, and *FPN2*_*prom*_∷*MtFPN2*-transformed *fpn2-1* plants. Bars = 500 μm. (D) Number of pink or white nodules in 28 dpi WT, *fpn2-1*, and *FPN2*_*prom*_∷*MtFPN2*-transformed *fpn2-1* plants. Data are the mean ± SE of at least 7 independently transformed plants. (E) Fresh weight of WT, *fpn2-1*, and *FPN2*_*prom*_∷*MtFPN2*-transformed *fpn2-1* plants. Data are the mean ± SE of at least 7 independently transformed plants. (F) Nitrogenase activity in 28 dpi nodules from WT, *fpn2-1*, and *FPN2*_*prom*_∷*MtFPN2*-transformed *fpn2-1* plants. Acetylene reduction was measured in duplicate from two sets of four pooled plants. Data are the mean ± SE. (G) Iron content in roots, shoot, and nodules of 28 dpi WT, *fpn2-1*, and *FPN2*_*prom*_∷*MtFPN2*-transformed *fpn2-1* plants. Data are the mean ± SE of 3 sets of 4 pooled plants. * indicates statistically significant differences (p < 0.05).

Consistent with the phenotype arising from altered iron delivery to the nodules, synchrotron-based micro-X-ray fluorescence studies of wild-type and *fpn2-1* nodules showed altered distribution of this element (Figure 5A). *MtFPN2* mutation led to less iron being allocated to the early fixation zones of the nodules, in those cases where the region was marginally developed (pink *fpn2-1* nodules). In the case of fully white nodules, no iron distribution indicative of ordered symbiosomes (iron-rich ring-like structures) was observed, consistent with no functional zone III being fully developed. This alteration in iron distribution had an impact on iron speciation as determined by micro-X-ray Absorption Near-Edge Spectroscopy (μXANES) on nodule sections. Different iron spectra were obtained from wild-type and mutant nodules (Figure 5B). Principal component analyses of the obtained spectra in the infection/differentiation zone, fixation zone and vascular region showed great iron speciation variation between wild-type and white *fpn2-1* nodules (Figure 5B). Interestingly, iron speciation in pink *fpn2-1* nodules showed an intermediate state midway from wild-type and white *fpn2-1* nodules. Although the complex iron speciation in nodules and the lack of a suitable number of spectra from many of these forms precludes a detailed description of the precise chemical species present in these organs, it was interesting to notice that the proportion of iron coordinated by sulfur (Fe-S clusters, including the iron-molybdenum cofactor of nitrogenase) was different in wild type, pink *fpn2-1*, and white *fpn2-1* nodules (Supplemental Table 1).

**Figure 5.**
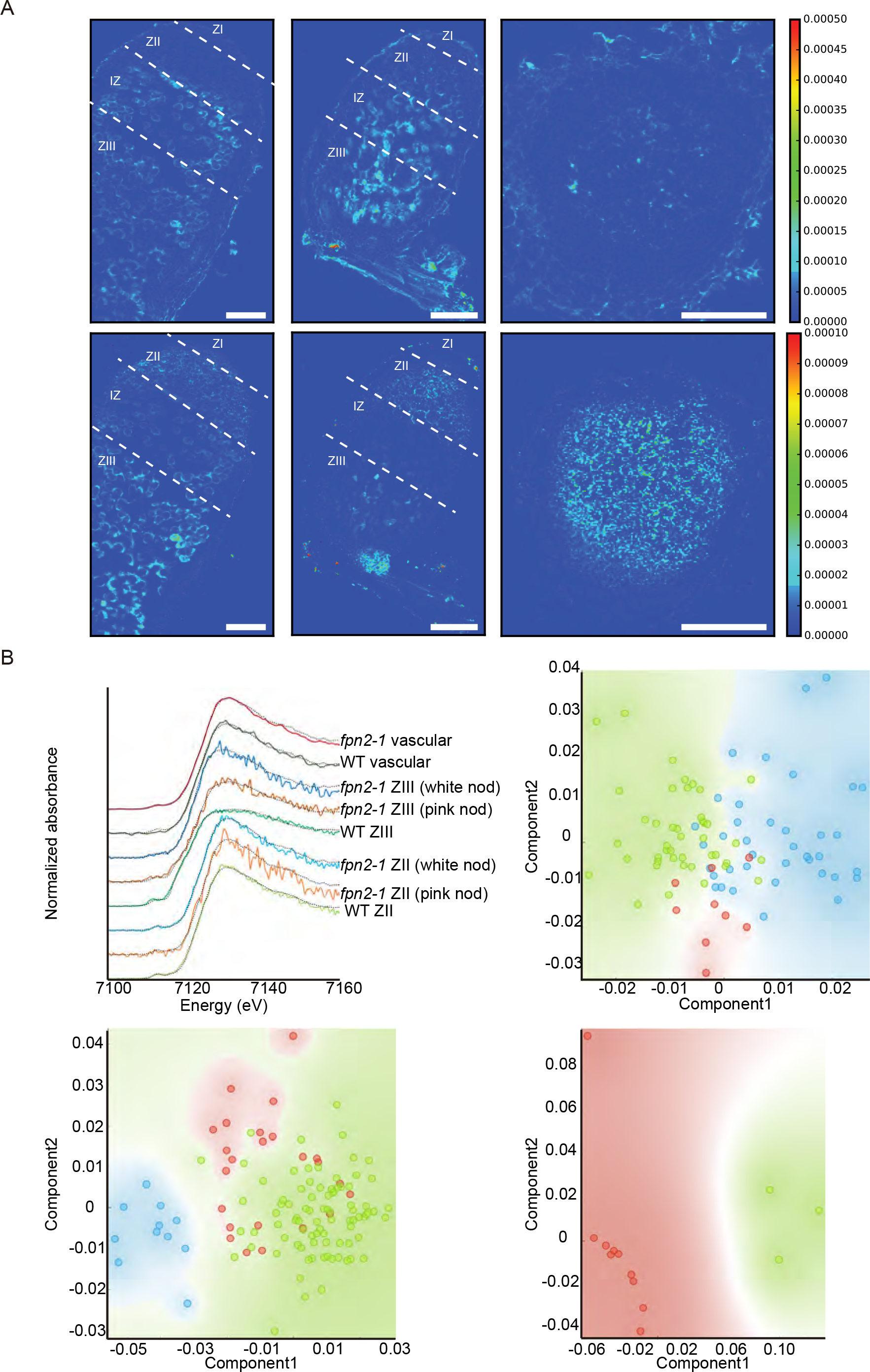
*MtFPN2* loss-of-function alters iron distribution and speciation in 28 dpi nodules. (A). Synchrotron-based X-ray fluorescence images of wild type (left), pink *fpn2-1* (middle), and white *fpn2-1* (right) nodules, showing calcium (top panels) or iron (lower panels) distribution. Bars = 200 μm, intensity scale in mass fraction units (g/g). (B) XANES spectra obtained from different regions of wild-type, pink *fpn2-1*, and white *fpn2-1* nodules (top left panel). This signal was decomposed into its two principal components for Zone II (top right panel), Zone III (lower left panel), and the vasculature (lower right panel). Green represents wild-type samples; red, pink *fpn2-1*; and blue, white *fpn2-1* nodules.

Considering the dual localization of MtFPN2 in the vasculature and in the symbiosomes, the reported phenotypes could be due to either iron not being released from the vessels and/or from it not getting across the symbiosome membrane. In an attempt to separate both causes, *MtFPN2* coding sequence was cloned under the *MtMOT1.3* promoter, a nodule-specific gene that we had previously located in the interzone and early fixation zone (Tejada-Jiménez et al., 2017). Mutants *fpn2-1* were transformed with this construct to test whether expressing *MtFPN2* only in the nodule cortex was sufficient to restore the phenotype. As shown in Figure 6, *MOT1.3*_*prom*_ driven expression of *MtFPN2* resulted in a partial improvement of plant growth (Figure 6A), with the development of functional pink nodules in significantly higher numbers than *fpn2-1* (Figures 6B and C). Overall, these plants showed improved biomass production compared to *fpn2-1* (Figure 6D), albeit not enough to reach wild-type levels. The improved iron delivery to *fpn2-1* led to significantly higher nitrogen fixation rates per nodule (Figure 6E). In contrast, when *MtFPN2* expression was driven by the promoter of vascular gene *MtNOOT1* (Magne et al., 2018), no significant differences were observed in terms of biomass production, nodule development, or nitrogenase activity with those that do not express it at all (Supplemental Figure 8).

**Figure 6.**
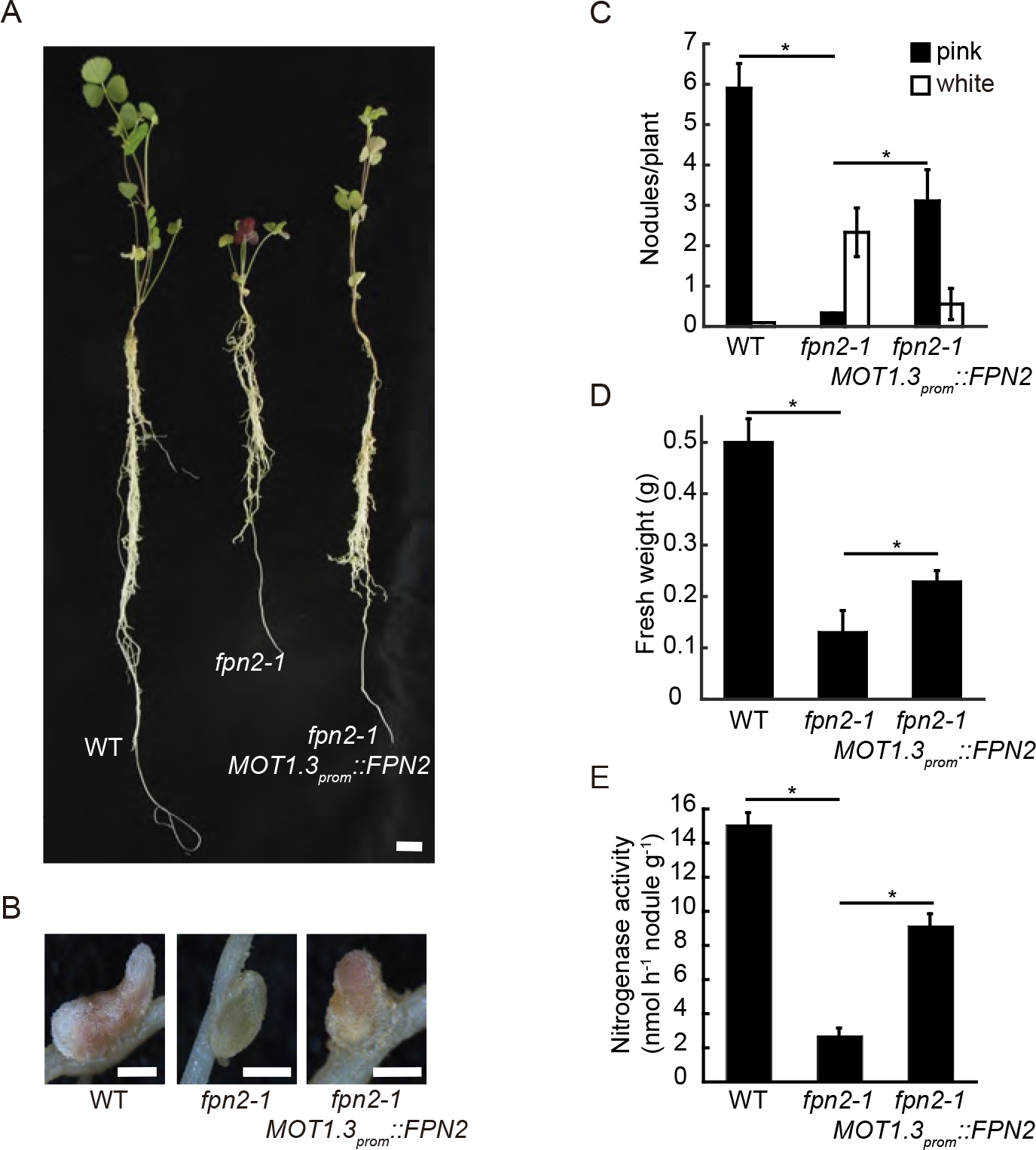
*MtFPN2* expression in cortical nodule cells partially restores the *fpn2-1* phenotype. (A) Growth of representative wild type (WT), *fpn2-1*, and *fpn2-1* transformed with the *MtFPN2* coding sequence expressed under the *MtMOT1.3* promoter (*MOT1.3*_*prom*_∷*MtFPN2*). Bar = 1 cm. (B) Detail of representative nodules of WT, *fpn2-1*, and *MOT1.3*_*prom*_∷*MtFPN2*-transformed *fpn2-1* plants. Bars = 500 μm. (C) Number of pink or white nodules in 28 dpi WT, *fpn2-1*, and *MOT1.3*_*prom*_∷*MtFPN2*-transformed *fpn2-1* plants. Data are the mean ± SE of at least 11 independently transformed plants. (D) Fresh weight of WT, *fpn2-1*, and *MOT1.3*_*prom*_∷*MtFPN2*-transformed *fpn2-1* plants. Data are the mean ± SE of at least 11 independently transformed plants. (E) Nitrogenase activity in 28 dpi nodules from WT, *fpn2-1*, and *MOT1.3*_*prom*_∷*MtFPN2*-transformed *fpn2-1* plants. Acetylene reduction was measured in duplicate from two sets of four pooled plants. Data are the mean ± SE. * indicates statistically significant differences (p < 0.05).

## DISCUSSION

Symbiotic nitrogen fixation carried out by the legume-rhizobia systems is one of the main sources of assimilable nitrogen in natural ecosystems and in agrosystems (Vance, 2001; Herridge et al., 2008). A key element in crop rotation strategies to increase nitrogen content in soils, symbiotic nitrogen fixation can minimize the use of polluting and expensive nitrogen fertilizers. Unveiling the molecular bases of this process is critical to improve nitrogen fixation rates and to introduce these capabilities to non-legume crops (Oldroyd and Dixon, 2014; Mus et al., 2016). However, for symbiotic nitrogen fixation to be efficiently used in sustainable agriculture or to extend it to other systems, close attention has to be paid to the mechanisms of delivery of essential, limiting nutrients, such as iron.

Nitrogen fixation, oxygen homeostasis, reactive oxygen species production and control, and several other nodule physiological processes rely on iron (Brear et al., 2013; González-Guerrero et al., 2014; González-Guerrero et al., 2016), a limiting nutrient in large areas of the world (Hirschi, 2009; White and Broadley, 2009). Therefore, nodules must ensure a steady supply of this element. While free-living diazotrophs can obtain iron directly from soil, endosymbiotic ones must obtain it from the host (Rodríguez-Haas et al., 2013; Udvardi and Poole, 2013). This process was likely adapted from the pre-existing long-distance iron transport systems. Consequently, we might expect similar mechanisms to those dedicated for iron delivery to the shoots, with the added handicap of the nodules becoming a second sink of an already scarce nutrient. Further supporting this hypothesis is the existence of a nodule-specific iron-regulated citrate efflux protein that showcases the importance of iron-citrate complexes in apoplastic iron solubility and in iron supply to symbiosomes (Kryvoruchko et al., 2018). Since ferroportins were associated to citrate-mediated long-distance iron supply (Morrissey et al., 2009), we speculated on a putative role for these proteins in symbiotic nitrogen fixation.

Model legume *M. truncatula* encodes three ferroportin-like proteins. One of them, MtFPN1, closely resembles the Mar1 protein located in the chloroplasts of Arabidopsis, that plays an undefined role in iron homeostasis in plants, as well as serving as entry-point for antibiotics (Conte et al., 2009). Similarly to *Mar1*, *MtFPN1* was expressed in the whole plant. The other two are orthologues to *A. thaliana* FPN1 and FPN2. While *MtFPN3* was expressed in most plant organs, *MtFPN2* transcripts were only detected in two discrete nodule regions: associated to the nodule vasculature, and from late Zone II to early Zone III. This dual localization indicates a dual function related with iron efflux from the cytosol, as inferred from the yeast complementation assays and from the available information on the biochemistry of this family of transporters.

In the vasculature, MtFPN2 would introduce iron into the lumen of an endomembranous compartment. Considering the relatively large amount of iron that has to be transferred across the nodule vessels into the nodules (González-Guerrero et al., 2014; González-Guerrero et al., 2016), it could be argued that the endomembranes would serve as an iron buffer, storing it prior to being delivered to the apoplast of the infection zone or to be recycled back to the host in older parts of the nodule. It could also be hypothesized that the iron concentrated in intracellular compartments could be released in the apoplast by exocytosis, as it has been proposed for phosphate transport mediated by the PHO1 protein (Arpat et al., 2012). It is unlikely that MtFPN2 would participate in a direct translocation of iron from the cytosol across the plasma membrane, since no signal was observed using immunolocalization and transmission electron microscopy. In contrast, the localization of MtFPN2 in the symbiosome provides a clear insight into a key function for MtFPN2: mediating iron delivery to the bacteroids.

Several evidences in addition to the expression and localization data support this role. Loss of *MtFPN2* function leads to a reduction of nitrogen fixation capabilities, with negative effects on plant growth, biomass production, and nodule development. Nitrogenase activity is highly dependent on iron delivery, due to its large requirements of iron for its three cofactors (Rubio and Ludden, 2005). Altering iron uptake by rhizobia-infected nodule cells presented a similar phenotype, as shown when characterizing the *nramp1-1 M. truncatula* mutant (Tejada-Jiménez et al., 2015). In addition, iron delivery to *fpn2-1* nodules is affected in terms of total concentration, distribution in the nodule, and speciation, indicating a large disturbance in nodule iron homeostasis. Altered nodule development has also been reported when legumes were grown in iron-deficient substrates (O’Hara et al., 1988; Tang et al., 1990; Tang et al., 1992), or when iron delivery was altered by mutating iron transporters (Hakoyama et al., 2012; Tejada-Jiménez et al., 2015), or the iron-chelation system (Stephens et al., 2011; Kryvoruchko et al., 2018). The changes in iron speciation in *fpn2-1* nodules would be the consequence of several factors: i) iron being stored or retained by iron-detoxification systems as a consequence of not reaching the proper acceptors, ii) loss of signal from ferro-proteins that have not received their cofactor, iii) mismetallation of proteins due to iron being retained in different compartments. Although attempts were made to fit the spectra obtained with different iron-complex reference standards, the complexity and number of these species made it extremely difficult to conclude anything other than a reduction of iron-sulfur species in nodules, which would very likely include the iron-sulfur clusters of nitrogenase. This phenotype was stronger in white non-functional nodules, but also present in those with wild type colour, a leaky phenotype that was also observed with *MtNramp1* mutants (Tejada-Jiménez et al., 2015) and that might indicated the existence of secondary systems able to deliver, with low affinity or non-specifically, some iron for nitrogen fixation in nodules.

The roles of MtFPN2 in the vasculature and in the symbiosomes were not equally important. Expressing *MtFPN2* only in the nodule cortical cells was enough to at least partially restore the wild-type phenotype. This means that facilitating iron transport across the symbiosome membrane would be the key function of MtFPN2. In contrast, when *MtFPN2* was only expressed in the vasculature, no significant improvements in plant growth, nodule development, or nitrogenase activity were observed compared to the *fpn2-1* nodules. This would also hint at the existence of mechanisms to deliver iron out of the vessels that are independent of MtFPN2. In contrast, in the symbiosomes, MtFPN2 would not be easily substituted by another iron transporter, including MtSEN1, which has also been hypothesized to be in this membrane (Hakoyama et al., 2012). This would also explain why the attempts to functionally complement *fpn2-1* by increasing iron content in the nutrient solution did not work, since it would require to: i) accumulate at high enough levels in the cell cytosol to overcome MtFPN2 absence without becoming toxic, and ii) recruit another transporter that might use iron as a substrate although at lower affinities.

Overall, we now have a better understanding of the mechanisms of iron delivery for symbiotic nitrogen fixation. Iron would be delivered by the vessels into the apoplast of the infection/differentiation zone of the nodule (Rodríguez-Haas et al., 2013). Iron release from the vessels would be facilitated by MtFPN2 and some other unknown protein(s). Once in the apoplast, it would form a complex with the citrate released by MtMATE67 (Kryvoruchko et al., 2018). Then, a yet-to-be-determined ferroreductase must work to convert Fe^3+^ into Fe^2+^ that is the substrate of MtNramp1, the protein responsible for iron uptake by rhizobia-infected nodule cells (Tejada-Jiménez et al., 2015). Once in the cytosol, iron will be sorted to different subcellular compartments. In the case of the symbiosome, MtMATE67 would also be required to form the iron-citrate complexes that seem to be the preferred iron species to be taken up by bacteroids (Moreau et al., 1995; Le Vier et al., 1996).

## METHODS

### Biological material and growth conditions

*M. truncatula* R108 and *fpn2-1* seeds were scarified in concentrated sulfuric acid for 7.5 min (R108) or 6.5 min (*fpn2-1*). Excess acid was removed and washed away eight times with cold water, followed by surface-sterilization in 50 % (v/v) bleach for 90 s. Sterilized seeds were embedded in sterile water overnight in darkness. Seeds were stratified in water-agar plates for 48 h at 4 °C and germinated at 22 °C. Seedlings were planted on sterile perlite pots, and inoculated with *S. meliloti* 2011 or the same bacterial strain transformed pHC60-GFP (Cheng and Walker, 1998). Plants were grown in a greenhouse under 16 h light / 8 h dark at 25 °C / 20 °C conditions. In the case of perlite pots, plants were watered every two days with Jenner’s solution or water alternatively (Brito et al., 1994). Nodulated plants were harvested at 28 dpi. For hairy-root transformation experiments, *M. truncatula* seedling were transformed with *Agrobacterium rhizogenes* strain ARqua1, containing the appropriate binary vector as described (Boisson-Dernier et al., 2001).

Yeast complementation assays were carried out using the *S. cerevisiae* strain *ccc1* MATa *ade2-1 his3-11 leu2-3,112, trp1-1, ura3-52 can1-100, ccc1∷his3*) and parental DY150 (MATa *ade2-1 his3-11 leu2-3,112, trp1-1, ura3-52 can1-100*). The strains were grown in synthetic dextrose (SD) medium, supplemented with the required auxotrophic nutrients, 2 % glucose as carbon source, and the iron concentration indicated in each particular plate.

### RNA Extraction and RT-qPCR

RNA was obtained from 28 dpi plants or equivalent age non-inoculated plants using Tri-Reagent (Life Technologies), treated with DNase turbo (Life Technologies), and cleaned with RNeasy Mini kit (Qiagen). Samples were tested for possible DNA contamination by PCR using the RNA samples as templates and primers specific for *M. truncatula Ubiquitin carboxyl-terminal hydrolase* (*Medtr4g077320.1*) (Supplemental Table 2). RNA integrity was confirmed by electrophoresis. cDNA was synthesized from 500 ng of DNA-free RNA using PrimeScript RT reagent Kit (Takara), supplemented with RNase out (Life Technologies).

Expression studies were carried out by real-time reverse transcription polymerase chain reaction (RT-qPCR) using the StepOne plus thermocycler (Applied Biosystems), using the Power SyBR Green master mix (Applied Biosystems). The primers used are indicated in Supplemental Table 2. RNA levels were standardized using the *M. truncatula Ubiquitin carboxyl-terminal hydrolase* (*Medtr4g077320.1*) gene. Real time cycler conditions have been previously described (González-Guerrero et al., 2010). The threshold cycle (Ct) was determined in triplicate from three different assays, each with at least pooled material from three different plants). The relative levels of expression were determined using the 2^ΔCt^ method. All assays contained a non-RT control to detect possible DNA contaminations.

### Yeast complementation assays

Restriction sites *PstI* and *XhoI* were added to *MtFPN2* cDNA by PCR using the primers listed in Supplemental Table 2. The amplicon was inserted into the *PstI*/*XhoI* sites of yeast expression vector pDR196 and cloned using T4 ligation. Yeast strains were transformed with either the empty pDR196 or pDR196∷*MtFPN2* using the lithium acetate method (Schiestl and Gietz, 1989) and selected by uracil autotrophy. Complementation assays were carried out in SD medium plates with the required auxotrophic nutrients. To test tolerance to iron, the plates were supplemented with 2.5 mM FeSO_4_.

### GUS Staining

*MtFPN2* promoter region was obtained by amplifying the 2144 pb upstream of the start codon using the primers Forward MtFPN2p-GW and Reverse MtFPN2p-GW (Supplemental Table 2). The amplicon was cloned in pDONR2017 (Invitrogen) by Gateway Technology (Invitrogen) and transferred to destination vector pGWB3 (Nakagawa *et al*., 2007). Hairy-root transformation of *M. truncatula* seedling was carried out as indicated above. After three weeks on Fahreus medium supplemented with 50 μg/ml kanamycin, the transformed plants were transferred to sterilized perlite pots and inoculated with *S. meliloti* 2011. GUS activity was determined in root and nodules of 28 dpi plants as described (Vernoud et al., 1999).

### Immunolocalization studies

The coding sequence region of *MtFPN2* and the promoter region used in the GUS staining were fused by PCR using the primers indicated in Supplemental Table 2. This fragment was cloned in pDONR207 (Invitrogen) using Gateway Technology and transferred to pGWB13 (three HA epitopes fused in C-terminus) (Nakagawa et al., 2007). Hairy-root transformation was carried out as described previously (Boisson-Dernier et al., 2001). Transformants were inoculated with *S. meliloti* 2011 containing the pHC60 plasmid that constitutively expressed GFP. Nodules and roots were collected at 28 dpi.

Samples for immunolocalization with a confocal microscope were fixed in 4% paraformaldehyde, 2.5 % sucrose in phosphate-buffered saline (PBS) solution at 4 °C overnight (12 – 16 h). Fixative was removed and samples washed twice. Nodule and roots were included in 6 % agarose and then sectioned in a Vibratome 1000 Plus (Vibratome) in 100 - 150 μm thick slices. Sections were dehydrated by serial incubation with methanol (30 %, 50 %, 70 % and 100 % in PBS) for 5 min and then rehydrated following the same methanol series in reverse order. Cell wall permeabilization was carried out by incubation with 2 % cellulase in PBS for 1 h and 0.1 % Tween 20 for 15 min. Sections were blocked with 5 % bovine serum albumin (BSA) in PBS and then incubated with 1:50 anti-HA mouse monoclonal antibody (Sigma) in PBS at room temperature for 2 h. Primary antibody was washed three times with PBS for 15 min and subsequently incubated with 1:40 Alexa 594 - conjugated anti-mouse rabbit monoclonal antibody (Sigma) in PBS at room temperature for 1 h. Secondary antibody was washed three times with PBS for 10 min, and then DNA was stained using DAPI. Images were obtained with a confocal laser-scanning microscope (Leica SP8) using excitation light at 488 nm and 561 nm for GFP and Alexa 594, respectively. DAPI-stained zones were visualized with UV light.

Immunolocalization in an electron microscope was carried out with 28 dpi nodules from MtFPN2-HA expressing plants fixed in 1 % formaldehyde, 0.5 % glutaraldehyde, 2.5% sucrose in 50 mM potassium phosphate buffer (pH 7.4) at 4 °C for 14 – 16 h in gentle agitation. Samples were dehydrated with ethanol series and embedded in LR-White resin (London Resin Company Ltd, UK). Nodule were placed in gelatine capsules, filled with LR-White resin and polymerized at 60 °C for 24 h. Serial ultrathin sections were cut using a diamond knife in a Reichert Ultracut S Ultramicrotrome (Leica). Sections for immunolocalization were blocked with 2 % BSA on PBS for 30 min, then incubated with 1:50 anti-HA rabbit monoclonal antibody (Sigma) in PBS at room temperature for 3 h. Primary antibody was washed ten times with PBS for 2 min and next incubated with 1:150 anti-rabbit goat monoclonal antibody conjugated to 15 nm gold particles (Fisher Scientific) in PBS at room temperature for 1 h. Secondary antibody was washed ten times with PBS for 2 min, after that samples were washed fifteen times with water for 2 min. Sections were stained with 2 % uranyl acetate and visualized in a JEM 1400 electron microscope at 80 kV at Centro Nacional de Microscopía Electrónica.

### Acetylene reduction assays

The acetylene reduction assay was used to measure nitrogenase activity in 28 dpi plants (Hardy et al., 1968). Wild type and *fpn2-1* roots with nodules were introduced in 30 ml vials fitted with rubber stoppers. Each tube contained four or five independently transformed plants. Three milliliters of air of each bottle was replaced by the same volume of acetylene. The vials were incubated for 30 min at room temperature, when 0.5 ml gas samples were taken from each of the tube. Ethylene content in each sample was measured in a Shimadzu GC-8A gas chromatograph using a Porapak N column. The amount of the ethylene produced was determined by measuring the height of the ethylene peak relative to standards.

### Metal content determination

Iron content was determined in shoots, roots, and nodules 28 dpi. Plant tissues were weighted and mineralized in 15.6 M HNO_3_ (trace metal grade) for 1 h at 80 °C and overnight at 20 °C. Digestions were completed with 2 M H_2_O_2_. Samples were diluted in 300 mM HNO_3_ prior to measurements. Element analyses were performed with Atomic Absorption Spectroscopy (AAS) in an AAnalyst 800 (Perkin Elmer), equipped with a graphite furnace. All samples were measured in duplicate.

### Synchrotron-based X-ray fluorescence and XANES

μXRF hyperspectral images and μXANES spectra were acquired on the beamline ID21 of the European Synchrotron Radiation Facility (Cotte et al., 2017), at 110 K in the liquid nitrogen (LN2) cooled cryostat of the Scanning X-ray Micro-spectroscopy end-station. A nodule from each of at least five *M. truncatula* R108 or *fpn2-1* plants (totalling five R108 nodules, five pink *fpn2-1* nodules, and five white *fpn2-1* nodules) were embedded into OCT medium and cryo-fixed by plunging in isopentane chilled with LN2. 25 μm-thick sections of frozen samples were obtained using a Leica LN22 cryo-microtome and accommodated in a Cu sample holder cooled with LN2, sandwiched between Ultralene (SPEX SamplePrep) foils. The beam was focused to 0.4×0.9 μm^2^ with a Kirkpatrick-Baez (KB) mirror system. The emitted fluorescence signal was detected with energy-dispersive, large area (80 mm^2^) SDD detector equipped with Be window (XFlash SGX from RaySpec). Images were acquired at the fixed energy of 7.2 keV, by raster-scanning the sample in the X-ray focal plane, with a step of 2×2 μm^2^ or 0.5×0.5 μm^2^ and 100 ms dwell time. Elemental mass fractions were calculated from fundamental parameters with the PyMca software package, applying pixel-by-pixel spectral deconvolution to hyperspectral maps normalized by the incoming current (Sole et al., 2007). The incoming flux was monitored using a drilled photodiode previously calibrated by varying the photon flux at 7.2 KeV obtaining a response of 1927.9 charges/photon with a linear response up to 200 kcps. In PyMCA the incoming flux and XRF detector parameters were set to 2×10^9^ photons/s, 0.7746 cm^2^ active area, and 4.65 cm sample to XRF detector distance. Sample matrix was assumed to be amorphous ice (11% H, 89% O, density 0.92 g/cm^3^), the sample thickness set at 25μm obtained with the use of a cryo-microtome.

Fe-K edge (7.050 to 7.165 keV energy range, 0.5 eV step) micro X-ray absorption near edge spectroscopy (μXANES) spectra were recorded in regions of interest of the fluorescence maps on ID21 beamline (ESRF, France). Individual spectra were processed using Orange software with the Spectroscopy add-on (Demsar et al., 2013). The pre-processing step consisted of vector normalization and a second derivative Savitsky-Golay filter to highlight potential differences among genotypes. Then a principal component analysis combined to a linear discriminant analysis (when there were more than two conditions to compare) was performed on the second derivative of the spectra to highlight potential differences among genotypes. Average spectra from the regions was then fitted using a linear combination routine

### Statistical tests

Data were analyzed by t-test to calculate statistical significance of observed differences, between wild type plants and analyzed mutants. Test results with p-values lower than 0.05 were considered as statistically significant.

## ACKNOWLEDGEMENTS

This research was funded by a Ministerio de Economía y Competitividad grant (AGL2015-65866-P), and a European Research Council Starting Grant (ERC-2013-StG-335284), to MGG. Development of *M. truncatula Tnt1* mutant population was, in part, funded by the National Science Foundation, USA (DBI-0703285) to KSM. The X-ray experiments were performed on beamline ID21 and ID16B at the European Synchrotron Radiation Facility (ESRF), Grenoble, France. We are grateful to Juan Reyes-Herrera at the ESRF for his assistance in using beamline ID21. We would also like to acknowledge the other members of laboratory 281 at Centro de Biotecnología y Genómica de Plantas (UPM-INIA) for their support and feedback in preparing this manuscript.

## AUTHOR CONTRIBUTION

VE carried out most of the experimental work. IA, CL, JV, and HC-M generated the synchrotron-based X-ray fluorescence, XANES, and X-ray tomographic data. JQ and JMA were responsible for metal determinations. MT-J and ER-N executed the yeast complementation assays. RP participated in *M. truncatula* transformation assays. JW and KSM performed *M. truncatula* mutant screening and isolated the *fpn2-1* allele. VE, JI, and MG-G were responsible for experimental design, data analyses, and wrote the manuscript.

