## Supplemental for "*Medicago truncatula* Ferroportin2 mediates iron import into nodule symbiosomes"

### SUPPLEMENTAL METHODS

#### Biological material and growth conditions

*Medicago truncatula* plants not inoculated with *S. meliloti* were watered every two weeks with a nutrient solution supplemented with 20 mM $NH_4NO_3$. Those that were supplemented with iron, were watered every week with 0.5 g/l of sequestrene.

#### Symbiosome isolation and MtFPN-GFP localization.

The coding sequence region of *MtFPN2* and the promoter region used in the GUS staining were fused by PCR using the primers indicated in Supplemental Table 2. This fragment was cloned in pDONR207 (Invitrogen) using Gateway Technology and transferred to pGWB4 (GFP-fused in C-terminus) (Nakagawa et al., 2007). Hairy-root transformation was carried out as described previously (Boisson-Dernier et al., 2001). Transformants were inoculated with a *S. meliloti* strain that expresses mCherry. Nodules and roots were collected at 28 dpi.

Symbiosome isolation for visualization of MtFPN2-GFP fusion was carried out from 28 dpi nodules homogenized in 2 % polyvinylpolypyrrolidone, 0.5 M sucrose, 50 mM Tris HCl pH 7.4, 10 mM DTT and 1 % protease inhibitor cocktail (Sigma), using the method by Catalano et al (Catalano et al., 2006). Symbiosomes were concentrated by centrifugation at 10,000 g for 1 min at 4 ºC, and resuspended in 0.5 M sucrose, 50 mM Tris HCl pH 7.4, 10 mM DTT and 1 % protease inhibitor cocktail. This solution was overlaid on another containing 1.5 M sucrose, 50 mM Tris-HCl pH 7.4, 10 mM DTT and 1 % protease inhibitor cocktail, centrifugated 5000 g for 30 s. The top phase and interphase were collected and concentrated by centrifugation at 10000 g for 90 s. The pellet was resuspended in 0.5 M sucrose, 50 mM Tris HCl pH 7.4, 10 mM DTT and 1 % protease inhibitor cocktail, and overlaid in 1 M sucrose, 50 mM Tris-HCl pH 7.4 and 10 mM DTT. Samples were centrifuged at 10.000 g for 5 min. The pellet was resuspended in 50 mM Tris-HCl pH 7.4, 10 mM DTT and 1 % protease inhibitor cocktail. Samples were observed in a Leica SP8 (Leica) confocal microscope using excitation light at 488 nm for GFP and 587 nm for mCherry.

#### Chlorophyll content

Chlorophyll was extracted from 50 mg of fresh leaves using 500 ml of dimethylformamide (DMF) at 4 °C for 24 h. Leaves were pelleted for 5 min at 600 g. A second extraction with 500 ml of DMF followed with strong agitation was carried out. The supernatant from both extractions were combined. Chlorophylls were measured spectrophotometrically at 647 and 664 nm, and concentrations determined as indicated by Inskeep and Bloom (1985).

**Nodule structure analyses**

Sections for nodule structural organization were obtained from 28 dpi nodules fixed in 1 % formaldehyde, 0.5 % glutaraldehyde, 2.5% sucrose in 50 mM potassium phosphate buffer (pH 7.4) at 4 °C for 14 – 16 h in gentle agitation. Samples were dehydrated with ethanol series and embedded in LR-White resin (London Resin Company Ltd, UK). Nodule were placed in gelatine capsules, filled with LR-White resin and polymerized at 60 °C for 24 h. Serial thin section of 1 µm, were cut using a diamond knife in a Reichert Ultracut S-Ultramicrotrome (Leyca), stained with a mixture of 1 % toluidine blue in aqueous 1 % Di-sodium tetra-borate in water Direct observation of sections was perfected by a Zeiss Axiophot photomicroscope (Carl Zeiss, Oberkochen, Germany) with pictures taken with a digital camera (Leica DFC 420C, Leica).

**X-ray tomography**

The 3D volumes have been acquired using X-ray nano-tomography (Cloetens et al., 1999) at the nano-analysis endstation ID16B at the ESRF having a focused beam of $50x50nm^2$ (Martínez-Criado et al., 2016). Phase contrast imaging was performed using this focused beam as secondary source producing a conic and monochromatic beam with an energy of 17.5 keV and a high flux of $6.10^{11}$ ph $s^{-1}$. Thanks to the geometry, the magnification i.e. pixel size and field of view depends of the sample position between the secondary source and the detector, in this experiment, pixel size of 323 nm has been used. For each tomography, about 1998 projections, were recorded on a PCO edge camera (2560×2160 pixels) along a 360° rotation with an exposure time of 100 ms per step. 3D reconstructions were achieved in two steps: *(i)* phase retrieval calculation using an inhouse developed octave script and *(ii)* filtered backprojection reconstruction using ESRF software PyHST2 (Mirone et al., 2014).

**Spectral fitting from XANES**

A reference library for Fe compounds was acquired on different material: Fe-foil (Fe(0)), Fe(II)-nicotianamine, Fe(II)S2 (http://ixs.iit.edu/database/), Fe(III)-haem (50 mM, pH7, bought from Sigma, CAS number: 16009-13-5), Fe(III)-cellulose (5 mM $FeCl_3$ + 50 mM cellulose, pH 5.8, bought from Sigma, CAS number: 9004-34-6), Fe(III) glutamic acid (5 mM $FeCl_3$ + 50 mM glutamic acid, pH 7, bought from Sigma, CAS number: 56-86-0) and Fe(III) ferritin (bought from Sigma, CAS number: 9007-73-2, 50 mM, pH7).

For linear combination fitting (LCF), reference compounds were classified as Fe(II)-S (FeS2), Fe(II)-O/N (Fe-NA) and Fe(III)-O (Fe-cellulose, Fe-glutamic acid and ferritin). Fe foil and Fe under haem form did not appear during the LCF process. XANES data treatment was performed using Athena software (Ravel and Newville, 2005) as previously described (Larue et al., 2014). Briefly, spectra were normalized and the reference compounds were used to perform LCF.

**Supplemental Figure 1.** Expression of *M. truncatula* ferroportin genes *MtFPN1* and *MtFPN3* in nodulated or nitrogen-fertilized *M. truncatula* plants. *Ubiquitin carboxyl-terminal hydrolase* (*MtUb1*) gene was used as a positive control for RT-PCRs.

**Supplemental Figure 2.** MtFPN2-GFP localizes in the symbiosomes. (A) MtFPN2-GFP distribution (green, left panel) in symbiosomes isolated from 28 dpi *M. truncatula* nodules inoculated with m-Cherry expressing *S. meliloti* (red, middle panel). Both images are overlaid with the transillumination file (right panel). Bars = 10 µm. (B) Negative control for GFP autofluorescence signal (right panel) obtained from 28 dpi *M. truncatula* nodules inoculated with m-Cherry expressing *S. meliloti* (red, middle panel). Both images are overlaid with the transillumination file (right panel). Bars = 10 µm.

**Supplemental Figure 3.** Controls for immunolocalization assays. (A) Longitudinal section of a 28 dpi *M. truncatula* nodule expressing *MtFPN2-HA* under its own promoter. No primary anti-HA antibody was used, but samples were incubated with secondary Alexa594-conjugated antibody (red, left panel). Transformed plants were inoculated with a GFP-expressing *S. meliloti* strain (green, middle panel). Both images were overlaid with the channel showing DAPI-stained DNA (blue, right panel). Bar = 100 µm. (B) Immunolocalization of MtFPN2-HA from 28 dpi *M. truncatula* nodule expressing *MtFPN2-HA* under its own promoter. No primary anti-HA antibody was used, but samples were incubated with secondary gold-conjugated antibody.

**Supplemental Figure 4.** MtFPN2-HA transports iron out of the cytosol. Parental yeast strain DY150 was transformed with the pDR196 empty vector, while the *ccc1* mutant was transformed with either pDR196 or pDR196 harboring the *MtFPN2* coding sequence fused to three C-terminal HA epitopes. Serial dilutions (10x) of the transformants were grown in YPD medium (top panel), SD medium with the required auxotrophic requirements (centre panel), or in SD medium with the required auxotrophic requirements and 2.5 mM $FeSO_4$ (lower panel).

**Supplemental Figure 5.** MtFPN2 is not required for plant growth under non-symbiotic conditions. (A) Growth of representative wild-type and *fpn2-1* plants when watered with a nutrient solution supplemented with ammonium nitrate and not inoculated with *S. meliloti*. Bar = 1 cm. (B) Fresh weight of wild-type and *fpn2-1* plants when watered with a nutrient solution supplemented with ammonium nitrate and not inoculated with *S. meliloti*. Data are the mean ± SE of at least 8 plants. (C) Chlorophyll content of wild-type and *fpn2-1* plants when watered with a nutrient solution supplemented with ammonium nitrate and not inoculated with *S. meliloti*. Data are the mean ± SE of at least 8 plants.

**Supplemental Figure 6.** Effect of *MtFPN2* mutation on nodule development. (A) Toluidine blue stain of longitudinal sections of 28 dpi wild-type, pink and white *fpn2-1* nodules. ZI indicates Zone I; ZII, Zone II; IZ, Interzone; and ZIII, Zone III. Bars = 200 µm. (B) Detail of Zone III cells from 28 dpi wild-type, pink and white *fpn2-1* nodules. Bars = 20 µm. (C) Longitudinal cross sections extracted from tomographic reconstructed volumes of 28 dpi wild type (left panel), pink and white *fpn2-1* nodules (centre and right panels, respectively). Bars = 100 µm

**Supplemental Figure 7.** Iron fortification of the nutrient solution does not rescue the *fpn2-1* phenotype. (A) Growth of representative 28 dpi wild-type and *fpn2-1* plants when watered with nutrient solution supplemented or not with sequestrene. Bar = 1 cm. (B) Fresh weight of 28 dpi wild-type and *fpn2-1* plants when watered with a nutrient solution supplemented or not with sequestrene. Data are the mean ± SE of at least 8 plants. (C) Nitrogenase activity in 28 dpi wild-type and *fpn2-1* nodules when watered with a nutrient solution supplemented or not with sequestrene. Acetylene reduction was measured in duplicate from two sets of four pooled plants. Data are the mean ± SE.

**Supplemental Figure 8.** *MtFPN2* expression in vascular nodule cells partially does not restore the *fpn2-1* phenotype. (A) Growth of representative wild type (WT), *fpn2-1*, and *fpn2-1* transformed with the *MtFPN2* coding sequence expressed under the *MtNOOT1* promoter ($NOOT1_{prom}$::*MtFPN2*). Bar = 1 cm. (B) Detail of representative nodules of WT, *fpn2-1*, and $NOOT1_{prom}$::*MtFPN2*-transformed *fpn2-1* plants. Bars = 500 µm. (C) Number of pink or white nodules in 28 dpi WT, *fpn2-1*, and $NOOT1_{prom}$::*MtFPN2*-transformed *fpn2-1* plants. Data are the mean ± SE of at least 11 independently transformed plants. (D) Fresh weight of WT, *fpn2-1*, and $NOOT1_{prom}$::*MtFPN2*-transformed *fpn2-1* plants. Data are the mean ± SE of at least 11 independently transformed plants. (E) Nitrogenase activity in 28 dpi nodules from WT, *fpn2-1*, and $NOOT1_{prom}$::*MtFPN2*-transformed *fpn2-1* plants. Acetylene reduction was measured in duplicate from two sets of four pooled plants. Data are the mean ± SE. * indicates statistically significant differences ($p < 0.05$).

**SUPPLEMENTAL FIGURE 1**

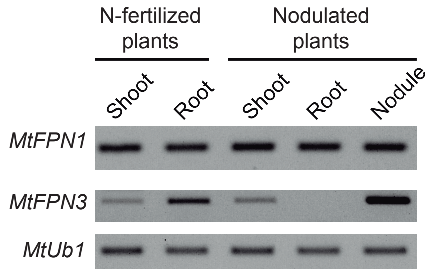


**Supplemental Figure 1.** Expression of *M. truncatula* ferroportin genes *MtFPN1* and *MtFPN3* in nodulated or nitrogen-fertilized *M. truncatula* plants. U*biquitin carboxyl-terminal hydrolase* (*MtUb1*) gene was used as a positive control for RT-PCRs.

**SUPPLEMENTAL FIGURE 2**

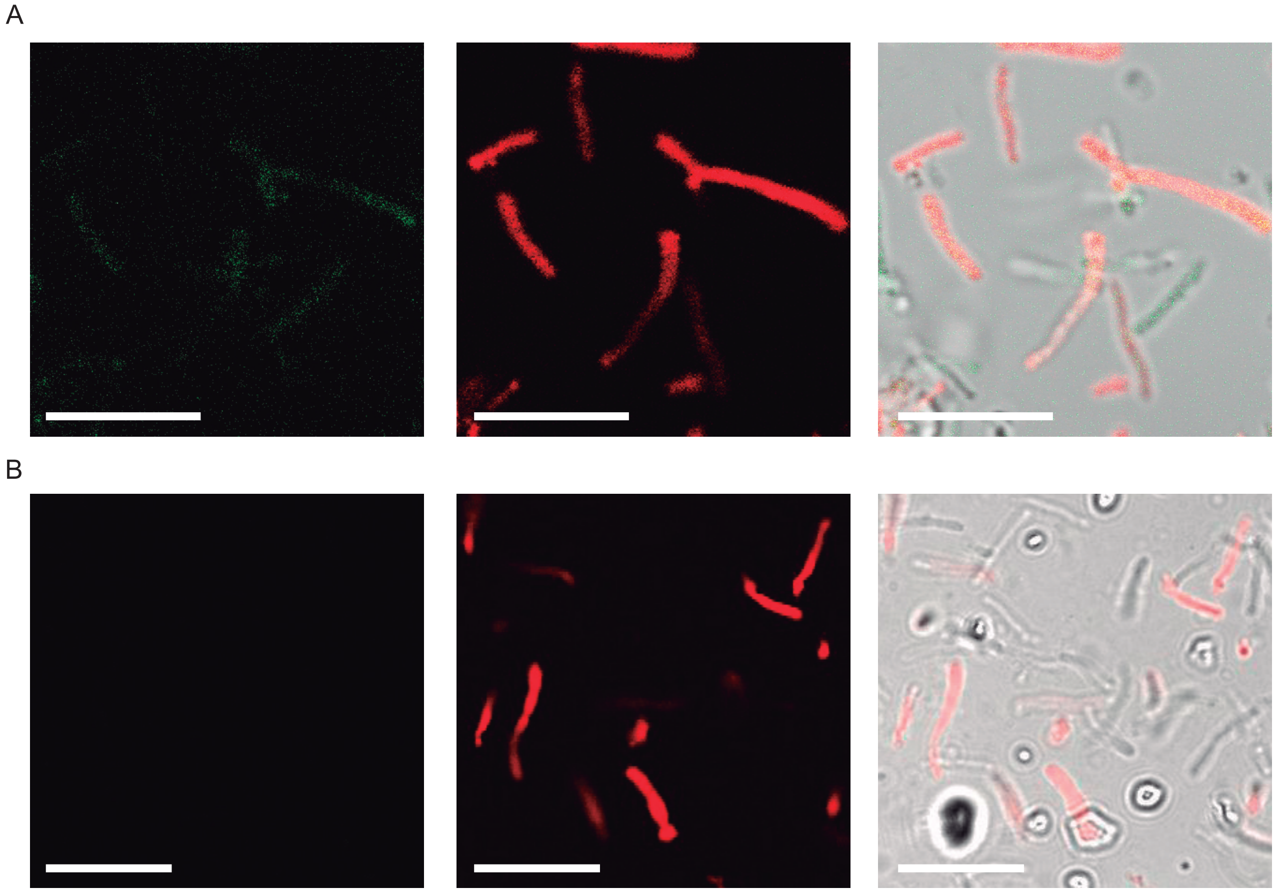


**Supplemental Figure 2.** MtFPN2-GFP localizes in the symbiosomes. (A) MtFPN2-GFP distribution (green, left panel) in symbiosomes isolated from 28 dpi *M. truncatula* nodules inoculated with m-Cherry expressing *S. meliloti* (red, middle panel). Both images are overlaid with the transillumination file (right panel). Bars = 10 μm. (B) Negative control for GFP autofluorescence signal (right panel) obtained from 28 dpi *M. truncatula* nodules inoculated with m-Cherry expressing *S. meliloti* (red, middle panel). Both images are overlaid with the transillumination file (right panel). Bars = 10 μm.

### SUPPLEMENTAL FIGURE 3

A

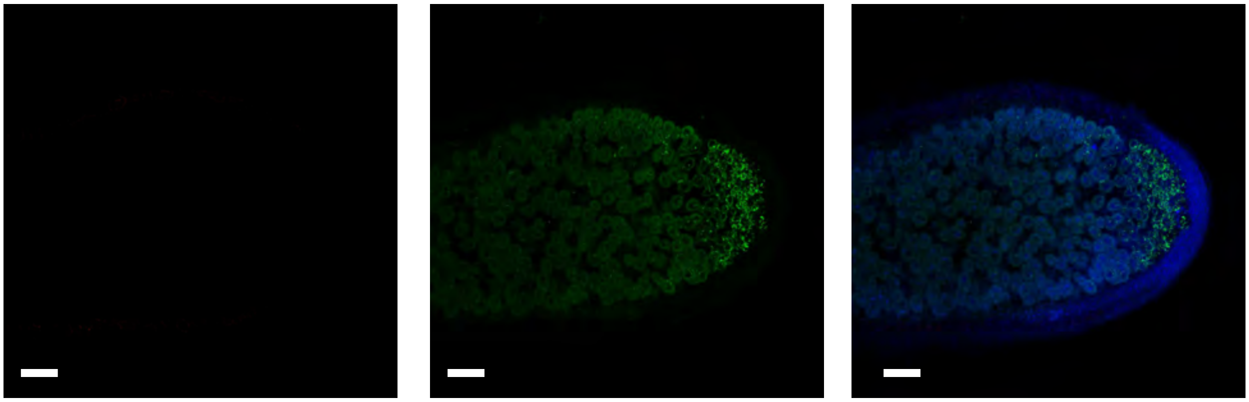

B

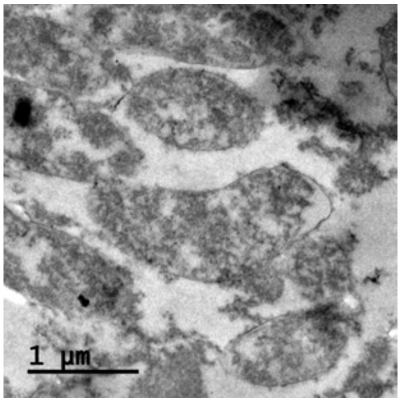


**Supplemental Figure 3.** Controls for immunolocalization assays. (A) Longitudinal section of a 28 dpi *M. truncatula* nodule expressing *MtFPN2-HA* under its own promoter. No primary anti-HA antibody was used, but samples were incubated with secondary Alexa594-conjugated antibody (red, left panel). Transformed plants were inoculated with a GFP-expressing *S. meliloti* strain (green, middle panel). Both images were overlaid with the channel showing DAPI-stained DNA (blue, right panel). Bar = 100 μm. (B) Immunolocalization of MtFPN2-HA from 28 dpi *M. truncatula* nodule expressing *MtFPN2-HA* under its own promoter. No primary anti-HA antibody was used, but samples were incubated with secondary gold-conjugated antibody.

**SUPPLEMENTAL FIGURE 4**

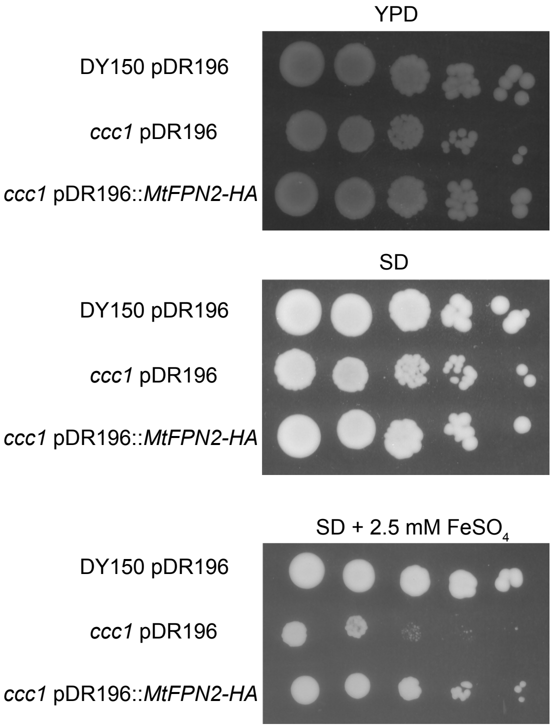


**Supplemental Figure 4.** MtFPN2-HA transports iron out of the cytosol. Parental yeast strain DY150 was transformed with the pDR196 empty vector, while the *ccc1* mutant was transformed with either pDR196 or pDR196 harboring the *MtFPN2* coding sequence fused to three C-terminal HA epitopes. Serial dilutions (10x) of the transformants were grown in YPD medium (top panel), SD medium with the required auxotrophic requirements (centre panel), or in SD medium with the required auxotrophic requirements and 2.5 mM $FeSO_4$ (lower panel).

**SUPPLEMENTAL FIGURE 5**

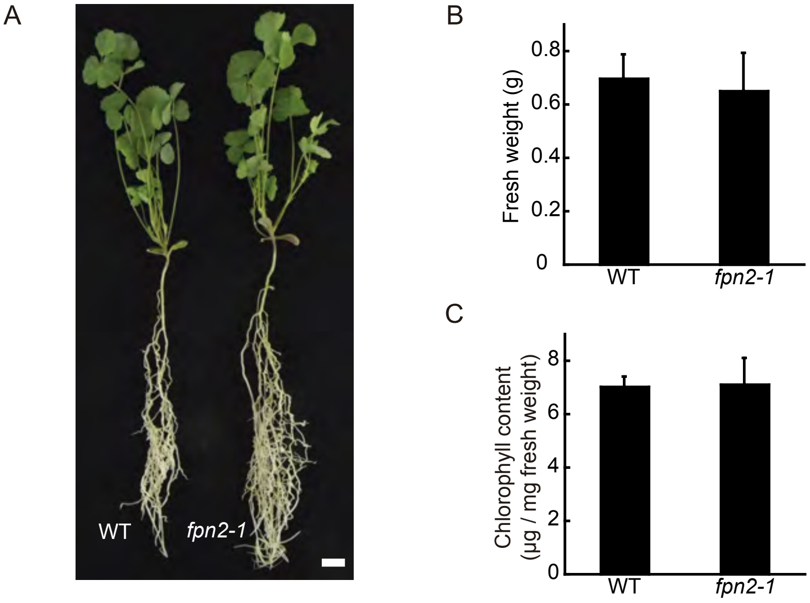


**Supplemental Figure 5.** MtFPN2 is not required for plant growth under non-symbiotic conditions. (A) Growth of representative wild-type and *fpn2-1* plants when watered with a nutrient solution supplemented with ammonium nitrate and not inoculated with *S. meliloti*. Bar = 1 cm. (B) Fresh weight of wild-type and *fpn2-1* plants when watered with a nutrient solution supplemented with ammonium nitrate and not inoculated with *S. meliloti*. Data are the mean ± SE of at least 8 plants. (C) Chlorophyll content of wild-type and *fpn2-1* plants when watered with a nutrient solution supplemented with ammonium nitrate and not inoculated with *S. meliloti.* Data are the mean ± SE of at least 8 plants..

### SUPPLEMENTAL FIGURE 6

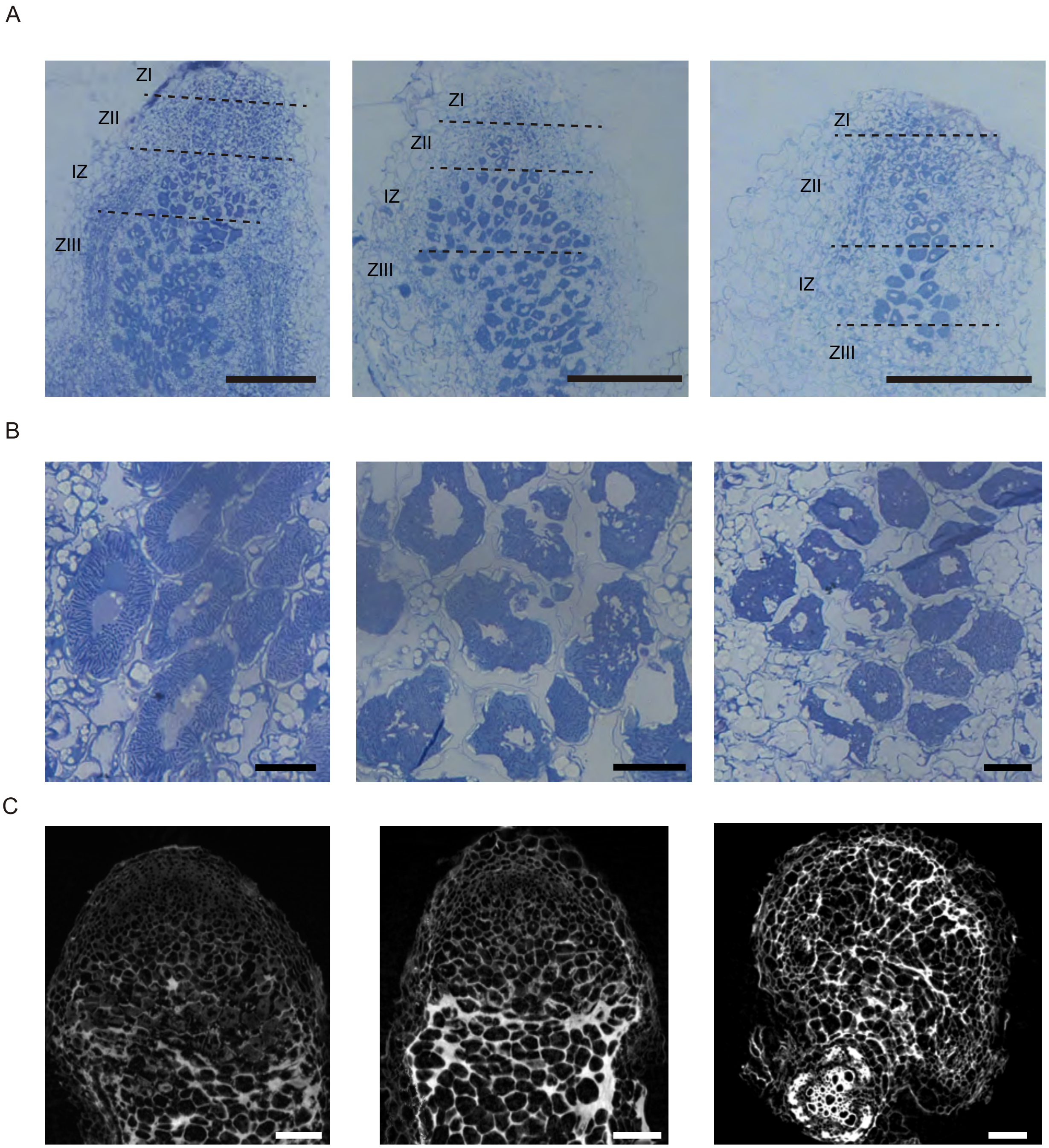


**Supplemental Figure 6.** Effect of *MtFPN2* mutation on nodule development. (A) Toluidine blue stain of longitudinal sections of 28 dpi wild-type, pink and white *fpn2-1* nodules. ZI indicates Zone I; ZII, Zone II; IZ, Interzone; and ZIII, Zone III. Bars = 200 µm. (B) Detail of Zone III cells from 28 dpi wild-type, pink and white *fpn2-1* nodules. Bars = 20 µm. (C) Longitudinal cross sections extracted from tomographic reconstructed volumes of 28 dpi wild type (left panel), pink and white fpn2-1 nodules (centre and right panels, respectively). Bars = 100 µm.

**SUPPLEMENTAL FIGURE 7**

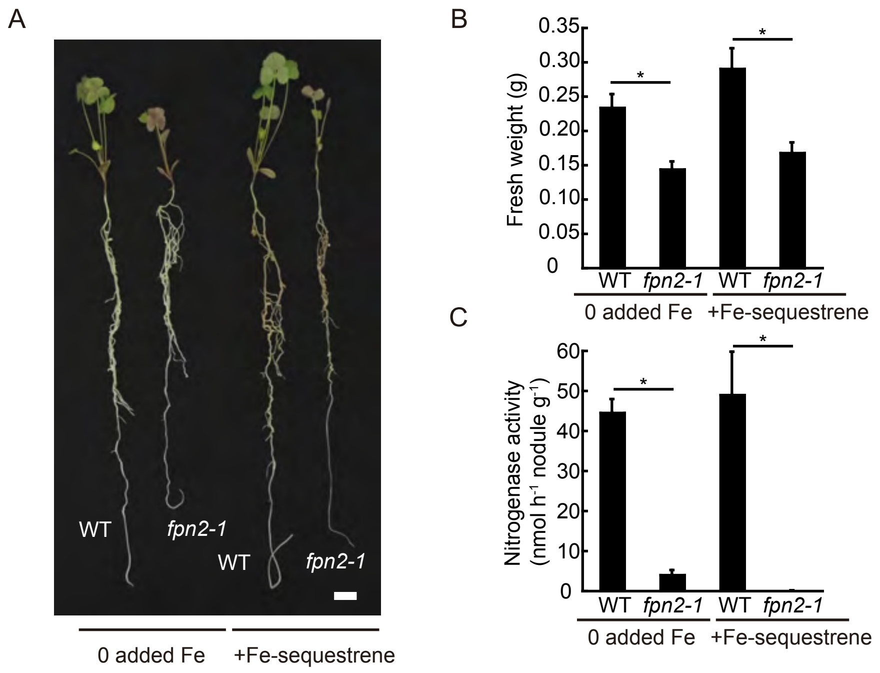


**Supplemental Figure 7.** Iron fortification of the nutrient solution does not rescue the *fpn2-1* phenotype. (A) Growth of representative 28 dpi wild-type and *fpn2-1* plants when watered with nutrient solution supplemented or not with sequestrene. Bar = 1 cm. (B) Fresh weight of 28 dpi wild-type and *fpn2-1* plants when watered with a nutrient solution supplemented or not with sequestrene. Data are the mean ± SE of at least 8 plants. (C) Nitrogenase activity in 28 dpi wild-type and *fpn2-1* nodules when watered with a nutrient solution supplemented or not with sequestrene. Acetylene reduction was measured in duplicate from two sets of four pooled plants. Data are the mean ± SE.

### SUPPLEMENTAL FIGURE 8

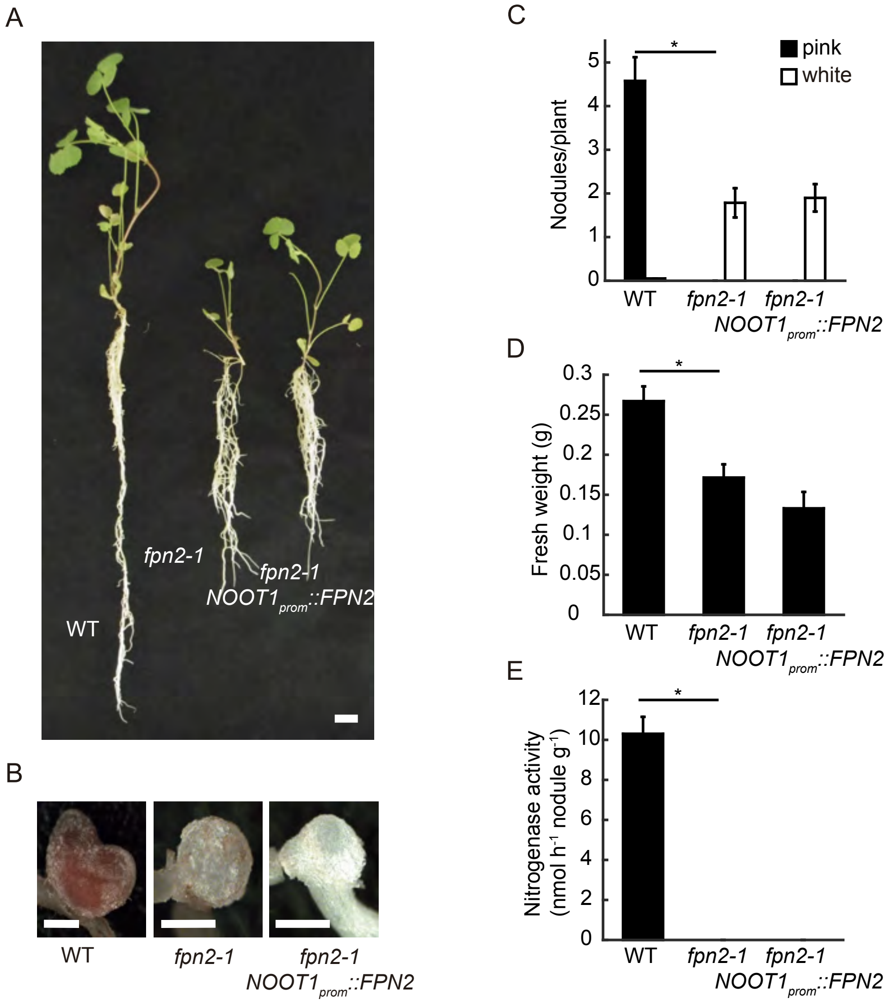


**Supplemental Figure 8.** *MtFPN2* expression in vascular nodule cells partially does not restore the *fpn2-1* phenotype. (A) Growth of representative wild type (WT), *fpn2-1*, and *fpn2-1* transformed with the *MtFPN2* coding sequence expressed under the *MtNOOT1* promoter ($NOOT1_{prom}$*::MtFPN2*). Bar = 1 cm. (B) Detail of representative nodules of WT, *fpn2-1*, and $NOOT1_{prom}$*::MtFPN2*-transformed *fpn2-1* plants. Bars = 500 µm. (C) Number of pink or white nodules in 28 dpi WT, *fpn2-1*, and $NOOT1_{prom}$*::MtFPN2*-transformed *fpn2-1* plants. Data are the mean ± SE of at least 11 independently transformed plants. (D) Fresh weight of WT, *fpn2-1*, and $NOOT1_{prom}$*::MtFPN2*-transformed *fpn2-1* plants. Data are the mean ± SE of at least 11 independently transformed plants. (E) Nitrogenase activity in 28 dpi nodules from WT, *fpn2-1*, and $NOOT1_{prom}$*::MtFPN2*-transformed fpn2-1 plants. Acetylene reduction was measured in duplicate from two sets of four pooled plants. Data are the mean ± SE. * indicates statistically significant differences ($p < 0.05$).

**Supplemental Table 1:** Iron speciation (%) in WT and *fpn2-1* nodules. n.d. = not detected, n.a. = not acquired (no region was present in the sectioned material).

| | WT | | | | *fpn2-1* (pink nodules) | | | | *fpn2-1* (white nodules) | | | |
|---|---|---|---|---|---|---|---|---|---|---|---|---|
| Region | Fe(II)-S | Fe(II)-O/N | Fe(III)-O | $R^2$ | Fe(II)-S | Fe(II)-O/N | Fe(III)-O | $R^2$ | Fe(II)-S | Fe(II)-O/N | Fe(III)-O | $R^2$ |
| ZII | 7 | 13 | 83 | 0.0005 | n.d. | 9 | 92 | 0.0089 | n.d. | 13 | 87 | 0.0015 |
| ZIII | 44 | 26 | 26 | 0.0055 | 26 | 22 | 52 | 0.0039 | 6 | 36 | 58 | 0.0036 |
| Vessels | n.d. | 21 | 78 | 0.0011 | 5 | 12 | 83 | 0.0005 | n.a. | n.a. | n.a. | - |

**Supplemental Table 2**: Primers used in this study

| Name | Sequence | Use |
|---|---|---|
| Fw MtFPN1 RT | CCCAAAGCCACACCACAAAG | RT-PCR of *MtFPN1* |
| Rv MtFPN1 RT | CCGTGCCTCCTACTTCAGTG | RT-PCR of *MtFPN1* |
| Fw MtFPN2 qRT | GGCTTGGACTGTGGATGTTTGACTTA | qRT-PCR of *MtFPN2* |
| Rv MtFPN2 qRT | ATAAACTCAACTTCCAGAACTCCTT | qRT-PCR of *MtFPN2* |
| Fw MtFPN3 RT | GGGATCACCTTTTTTTTTGGCTGAGA | RT-PCR of *MtFPN3* |
| Rv MtFPN3 RT | TTGGTAAGTCAACTTATCAACCCATGTTCC | RT-PCR of *MtFPN3* |
| Fw MtUBQ qRT | GAACTTGTTGCATGGGTCTTGA | qRT-PCR of *M. truncatula Ubiquitin carboxyl-terminal hydrolase* |
| Rv MtUBQ qRT | CATTAAGTTTGACAAAGAGAAAGAGACAGA | qRT-PCR of *M. truncatula Ubiquitin carboxyl-terminal hydrolase* |
| Fw MtFPN2p - GW | GGGGACAAGTTTGTACAAAAAAGCAGGCTTGA GACAGATAAGGCAGATA | Cloning of *MtFPN2* promoter in Gateway vector |
| Rv MtFPN2p -1GW | GGGGACCACTTTGTACAAGAAAGCTGGGTATT TATCATATGAACAAGAGT | Cloning of *MtFPN2* promoter in Gateway vector |
| Rv MtFPN2 - GW | GGGGACCACTTTGTACAAGAAAGCTGGGTTTG CAGAAGAATTAATCGACC | Cloning of *MtFPN2* gene in Gateway vector |
| Fw MtFPN2p - CDS FPN2 | ACTCTTGTTCATATGATAAAATGGATGAAAAA GAAGTTTT | Fusion PCR between *MtFPN2* promoter and *MtFPN2* coding sequence |
| Rv MtFPN2p - CDS FPN2 | AAAACTTCTTTTTCATCCATTTTATCATATGAA CAAGAGT | Fusion PCR between *MtFPN2* promoter and *MtFPN2* coding sequence |
| 5MtFPN2ATGPstI | TTTCTGCAGATGGATGAAAAAGAAGTTTTGAG AAAG | Cloning of *MtFPN2* coding sequence in pDR196 |
| 3MtFPN2STOPXhoI | TTTCTCGAGTCATGCAGAAGAATTAATCGACC AT | Cloning of *MtFPN2* coding sequence in pDR196 |
| Rv MtFPN2p - CDS FPN2 GW | GGGGACCACTTTGTACAAGAAAGCTGGGTTCA TGCAGAAGAATTAATCG | Cloning of *MtFPN2*$_{prom}$::*MtFPN2* cloning sequence in Gateway vector |
| Fw MtMOT1.3p-2000 GW | GGGGACAAGTTTGTACAAAAAAGCAGGCTAGT GCATAGACAACATCAAAAAGC | Fusion of *MtMOT1.3* promoter with *MtFPN2* and cloning in Gateway vector |
| Rv MtMOT1.3p-CDS FPN2 | CAAAACTTCTTTTTCATCCATATGGAATGTGAT GTGTTAATTGGC | Fusion of *MtMOT1.3* promoter with *MtFPN2* |
| Fw MtMOT1.3p CDS FPN2 | GCCAATTAACACATCACATTCCATATGGATGA AAAAGAAGTTTTG | Fusion of *MtMOT1.3* promoter with *MtFPN2* |
| Fw MtNOOT1p-2000 GW | GGGGACAAGTTTGTACAAAAAAGCAGGCTCTT TCCTTTCTTTCTTTCCA | Fusion of *MtNOOT1* promoter with *MtFPN2* and cloning in Gateway vector |
| Rv MtNOOT1p-CDS FPN2 | CAAAACTTCTTTTTCATCCATTACATCTTCAAT CGAGGCTTTTTC | Fusion of *MtNOOT1* promoter with *MtFPN2* |
| Fw MtNOOT1p CDS FPN2 | GAAAAAGCCTCGATTGAAGATGTAATGGATGA AAAAGAAGTTTTG | Fusion of *MtNOOT1* promoter with *MtFPN2* |
